# Determining the core bacterial and fungal genera in table olive fermentations

**DOI:** 10.1101/2025.03.16.643505

**Authors:** Annamaria Ricciardi, Francisco Noé Arroyo López, Marilisa Giavalisco, Rocchina Pietrafesa, Eugenio Parente

**Affiliations:** Dipartimento di Scienze Agrarie, Forestali, Alimentari ed Ambientali, Università degli Studi della Basilicata, Potenza, Italy; Food Biotechnology Department. Instituto de la Grasa, Consejo Superior de Investigaciones Cientificas (CSIC), Seville, Spain

**Keywords:** Table olives, core microbiota, fungal community, bacterial community, metataxonomic analysis

## Abstract

Table olives are among the most ancient and important fermented foods of the Mediterranean basin. Their production is still strongly related to traditional practices, and the lack of thermal treatments, the reliance on natural contamination and selective factors (NaCl, pH, occurrence of phenolics, etc.) determine the dynamics of the microbial community. Lactic acid bacteria (LAB) and yeasts have a pivotal role in table olive microbial communities, but several halophilic and alkalophilic microorganisms may also contribute, positively or negatively, to the quality and safety of this fermented vegetable.

We have use metataxonomic data extracted from the FoodMicrobionet database to provide quantitative insights on the structure of bacterial and fungal microbial communities of table olives and to identify core genera in different trade preparations.

*Celerinatantimonas* and *Lactiplantibacillus* were the most prevalent genera among bacteria, followed by several LAB, halophilic and alkalophilic lactic acid bacteria (HALAB) and Gram negatives, including non-halophilic species. Similarly, 3 fungal genera (*Pichia, Candida,* and *Wickerhamomyces*) were the most abundant and prevalent among fungi.

The distribution of both bacteria and fungi varied significantly in different olive varieties, among olives, brines and contact surfaces or materials, and at different production stages, and no clear grouping related to the combination of ripeness and trade preparation was found, although HALAB were characteristically abundant in Spanish style green olives. Addition of starter cultures affected the composition and dynamics of microbial communities to a variable extent.

**Highlights:**

- Data from 16 metataxonomic studies on table olives, including 833 samples, were combined.
- Enhanced metadata facilitated the comparison among different studies.
- Composition of the microbiota of table olives was highly variable.
- *Celerinatantimonas* and *Lactiplantibacillus* dominated the core bacterial microbiota.
- *Pichia*, *Candida,* and *Wickermahomyces* were the most prevalent among fungi.

## 1. Introduction

Table olives are the result of the processing fruits of selected varieties of *Olea europea* L. *sativa* that have been treated to remove their natural bitterness and preserved by fermentation, heat treatment, drying or salting (IOOC, 2004). The preservation of olives in brines dates back to 7,000 BP (Langgut and Garfinkel, 2022). Nowadays, this fermented vegetable is still produced mostly in the Mediterranean Basin, but the production and export from the Americas is increasing. Thereby, world production for the crop year 2023/24 is estimated at 2,828,500 t, and is concentrated in Mediterranean countries such as Spain, Egypt, Turkey, Morocco, Italy, and Greece (source: https://www.internationaloliveoil.org/world-market-of-olive-oil-and-table-olives-data-from-december-2024/, last accessed March 2025).

While the level of innovation in industrial production may be high (Campus et al., 2018), table olive production for artisanal producers is still strongly related to traditional practices, and, as for many other vegetable fermentations (Gänzle, 2022; Hernández-Velázquez et al., 2024; Tamang et al., 2020), the lack of thermal treatments of raw material, the reliance on natural contamination and selective factors (NaCl, acidic pH, occurrence of phenolics, etc.) determine the dynamics of the microbial community and could affect the quality and safety of the final product (Anagnostopoulos and Tsaltas, 2022; Perpetuini et al., 2020; Vaccalluzzo et al., 2020). On the other hand, several chemical and physical treatments are used at the time of packaging to enhance stability of commercial products (Campus et al., 2018; Panagou et al., 2003).

IOOC recognizes 5 main trade preparation groups for table olives: i) treated olives (often referred to as Spanish style olives, treated by alkali, prior to fermentation in brine); ii) natural olives (Greek style olives, naturally fermented in brine with no alkali treatment); iii) dehydrated and/or shrivelled olives (which may or may not be treated in brines); iv) olives darkened by oxidation (California style; which may be fermented prior to oxidation treatment); and v) other specialties (which do not fall in previous categories) (https://www.internationaloliveoil.org/wp-content/uploads/2019/11/COI-OT-NC1-2004-Eng.pdf). Microorganisms play an important role in the production, safety and quality of all these products (Anagnostopoulos and Tsaltas, 2022; Vaccalluzzo et al., 2020).

Several PDO and PGI table olive varieties are recognized in the European Union (https://ec.europa.eu/agriculture/eambrosia/geographical-indications-register/) and their microbiota may contribute to aspects related to identity and quality (Anagnostopoulos and Tsaltas, 2022). Additionally, the composition of the microbiota may be used to prevent frauds or identify the geographical origin of the product (Argyri et al., 2020; Kalogiouri et al., 2020; Kamilari et al., 2019).

Natural contamination from the olives, use of salt, spices, water, food contact surfaces and, in some cases, the addition of starter cultures, determine the assembly of the initial microbial community (Lanza, 2013; Penland et al., 2021), whose evolution is affected by a large number of factors, including initial pH, scarcity of nutrients (due to the repeated washing needed to remove alkali residuals in some varieties), high phenolic concentration, high salt concentration (brines with 6-12% NaCl are used, although there is a trend for reduction of salt concentration), temperature (which may be low in some areas at the beginning of fermentation), and anaerobiosis in large fermenters (Anagnostopoulos and Tsaltas, 2022; Benítez-Cabello et al., 2023; Blekas et al., 2002; Khalil et al., 2023; Perpetuini et al., 2018).

Due to the diversity of contamination sources, to the lack of heat treatments, and to the variety and variability of ecological factors influencing this fermentation process, the microbial communities of table olives are complex and dynamic.

As for many other food products, -omic approaches are increasingly being used to complement or even replace cultivation-based approaches (Anagnostopoulos and Tsaltas, 2022; Ferrocino et al., 2022; De Filippis et al., 2018; Tsoungos et al., 2023). Two main microbial groups, lactic acid bacteria (LAB) (Giavalisco et al., 2024; Portilha-Cunha et al., 2020) and yeasts (Arroyo-López et al., 2008, 2012), are recognized as having a pivotal role in table olive microbial communities, but several halophilic and alkalophilic microorganisms may also contribute, positively or negatively, to the quality and safety of table olives (de Castro et al., 2022; Lucena-Padrós and Ruiz-Barba, 2016; Posada-Izquierdo et al., 2021).

The microbiology of table olives has been reviewed multiple times, including recently (Arroyo-López et al., 2018; Anagnostopoulos and Tsaltas, 2022; Campus et al., 2018; Heperkan, 2013; Perpetuini et al., 2020; Portilha-Cunha et al., 2020; Tsoungos et al., 2023; Vaccalluzzo et al., 2020). These reviews provide qualitative data on the occurrence, dynamics and abundance of beneficial, functional, and spoilage microorganisms in table olive fermentation, but they usually lack quantitative insights nor provide re-usable data.

At the time of writing this article, there were 36 studies which investigated the microbiota of table olives using amplicon targeted approaches (Table 1). In most cases, studies focused on one or few olive varieties and trade preparations; while the most frequent approaches include targeting of regions the 16S rRNA gene or ITS region using the Illumina platform, the type of samples used (fruits, brines, or both; environmental samples) varies among studies. The quality of metadata is also highly variable, as are the experimental designs, although the comparison of started/unstarted fermentations or the evaluation of the effect of salt concentration is rather frequent. Qualitative lists of genera found in most studies have been presented in recent reviews (Tsoungos et al., 2023) but they are of little help in evaluating the variability in prevalence and abundance of the different species or in the identification of core microbial populations of fermented olives in general or of selected processes and no public repository with processed data on the microbiota of table olives is available. There is therefore a need for a reanalysis of published data in order to provide a comprehensive view of the structure and dynamics of the microbial communities of table olives and to foster access to this data by scientists and stakeholders.

**Table 1.** Metataxonomic studies on the microbial communities of table olives (in inverse chronological order).

| Olive varieties | Olive trade preparation | Platform | Targeted region(s) | Description | Reference |
| --- | --- | --- | --- | --- | --- |
| **Kalamata** | Greek style olives, black | Oxford Nanopore | 16S rRNA gene | Use of probiotic LAB starters in the production of Kalamata black olives | (Vougiouklaki et al., 2024) |
| **Manzanilla** | Spanish style olives, green | Illumina | ITS region | Impact of the addition of LAB starters on the fungal microbiota | (López-García et al., 2024) |
| **Gordal, Manzanilla, Hojiblanca** | Greek style olives, green | Illumina | 16S rRNA gene, ITS region | Impact of wild yeast starters on the volatilome | (Ruiz-Barba et al., 2024) |
| **Taggiasca** | Greek style olives, turning color | Illumina | 26S rRNA gene | Dynamics of yeast species during natural fermentation | (Traina et al., 2024) |
| **Thassos** | Naturally dry salted olives, black | Illumina | 26S rRNA gene | Composition of yeast microbiota in dry salted olives | (Gounari et al., 2023) |
| **Cypriot, Picual, Kalamata** | Greek style, green or black | Illumina | 16S rRNA gene, ITS region | Bacterial and fungal microbiota of three varieties, collected in different locations | (Kamilari et al., 2023) |
| **Halkidikis, Thassou, Kalamon, Amfissis, and Konservolia** | Greek style, Spanish style, dry salted | Illumina | 16S rRNA gene, 18S rRNA gene | Composition of bacterial and yeast microbiota in market samples of dry olives | (Mougiou et al., 2023) |
| **Manzanilla** | Spanish style, green | Illumina | 16S rRNA gene, ITS region | Fungal and bacterial microbiota in laboratory fermented olives inoculated with LAB and yeasts | (López-García et al., 2023) |
| **Manzanilla** | Spanish or Greek style, green | Illumina | 16S rRNA gene, ITS region | Bacterial and yeast microbiota in uninoculated olive fermentation | (Ruiz-Barba et al., 2023a) |
| **Gordal, Hojiblanca and Manzanilla** | Greek style, green | Illumina | 16S rRNA gene, ITS region | Bacterial and yeast microbiota in uninoculated natural green olive fermentation | (Ruiz-Barba et al., 2023b) |
| **Manzanilla** | Spanish style, green | Illumina | 16S rRNA gene, ITS region | Optimization of a pretreatment technique for recovery of LAB and yeasts from biofilms | (López-García et al., 2022) |
| **Manzanilla, Aloreña, Hojiblanca, Gordal, Morona, Verdial, Empeltre, Empeltre Mallorquina, Kalamata, Conservolea, Galega, Picholine, Criolla, Lucques du Languedoc, Azapa, Arauco** | Various | Illumina | ITS region | Fungal communities of packaged olives | (Benítez-Cabello et al., 2022) |
| **Nyons** | Greek style, black | Illumina | 16S rRNA gene, ITS region | Effect of salt reduction and inoculation with a complex consortium or backslipping on microbiota dynamics | (Penland et al., 2022) |
| **Hojiblanca** | Spanish style, green | Illumina | 16S rRNA gene | Effect of inoculation with LAB on the dynamics of bacterial microbiota | (Correa-Galeote et al., 2022) |
| **Gordal, Hojiblanca and Manzanilla** | Spanish style, green | Illumina | 16S rRNA gene | Investigation of the role of *Celerinatantimonas* in gas pocket defect | (de Castro et al., 2022) |
| **Ascolana tenera** | Spanish and Greek style, green | Illumina | 16S rRNA gene | Impact of starter addition on the microbiota in laboratory scale fermentation | (Maoloni et al., 2022) |
| **Halkidiki, Conservolea** | Spanish and Greek style, green | Illumina | ITS region | Fungal microbiota of green olives stored in two types of pouches | (Tzamourani et al., 2022) |
| **Nyons** | Greek style, black | Illumina | 16S rRNA gene, ITS region | Source tracking of desirable and spoilage microorganisms | (Penland et al., 2021) |
| **Aloréña de Malaga** | Greek style, green, cracked | Illumina | 16S rRNA gene, ITS region | Microbiota of traditionally packed olives during storage | (López-García et al., 2021) |
| **Kalamata** | Greek style, black | Illumina | 16S rRNA gene, 18S rRNA gene | Evolution of microbiota during storage in modified atmosphere | (Michailidou et al., 2021) |
| **Unknown/Unspecified** | Greek style, green, cracked | Illumina | Unknown | Composition of the microbiota fungal and bacterial microbiota in brine of an unspecified Turkish variety | (Demirci et al., 2021) |
| **Manzanilla** | Spanish style, green | Illumina | 16S rRNA gene | Diversity of the bacterial microbiota in spoiled industrial fermentations | (Arroyo-López et al., 2021) |
| **Manzanilla** | Spanish style, green | Illumina | 16S rRNA gene | Impact of a LAB and yeast starter on bacterial communities | (Benítez-Cabello et al., 2020a) |
| **Kalamata** | Greek style, black | Illumina | 16S rRNA gene, ITS region | Bacterial and fungal microbiota of olives and brines of industrially fermented Kalamata olives | (Kazou et al., 2020) |
| **Nyons** | Greek style, black | Illumina | 16S rRNA gene, ITS region | Dynamics of fungal and bacterial microbiota during fermentation | (Penland et al., 2020) |
| **Picual** | Greek style, green | Illumina | 16S rRNA gene | Dynamics of bacterial community in uninoculated and inoculated (LAB) fermentation | (Anagnostopoulos et al., 2020) |
| **Konservolia and Halkidiki** | Greek style, black and Spanish style, green | Ion Torrent (bacteria), Illumina (fungi) | 16S rRNA gene, ITS region | Bacterial and fungal communities in samples from different regions | (Argyri et al., 2020) |
| **Manzanilla, Aloreña, Hojiblanca, Gordal, Morona, Verdial, Empeltre, Empeltre Mallorquina, Kalamata, Conservolea, Galega, Picholine, Criolla, Lucques du Languedoc, Azapa, Arauco** | Various | Illumina | 16S rRNA gene | Bacterial communities of commercial packaged olives | (Benítez-Cabello et al., 2020b) |
| **Manzanilla, Hojiblanca** | California style | Illumina | 16S rRNA gene | Bacterial communities at different stages of processing | (Medina et al., 2018) |
| **Manzanilla, Gordal** | Spanish style, green | Illumina | 16S rRNA gene, ITS region | Bacterial and fungal microbiota of spoiled olives | (de Castro et al., 2018) |
| **Nocellara Etnea** | Greek style, green | Ion Torrent | 16S rRNA gene | Bacterial microbiota dynamics in LAB started and spontaneous fermentation | (Randazzo et al., 2017) |
| **Aloreña de Malaga** | Greek style, green | Illumina | 16S rRNA gene | Bacterial microbiota in cured or cracked olives, with or without heat shock | (Rodríguez-Gómez et al., 2017) |
| **Nocellara del Bélice** | Spanish style and Picholine, green | Illumina | 16S rRNA gene | Effect of salt reduction and type of processing on bacterial microbiota | (Zinno et al., 2017) |
| **Aloreña de Malaga** | Greek style, green | F454 | ITS region | Dynamics of fungal microbiota in brines and fruit | (Arroyo-López et al., 2016) |
| **Aloreña de Malaga** | Greek style, green | F454 | 16S rRNA gene | Dynamics of bacterial microbiota in brines and fruit | (Medina et al., 2016) |
| **Nocellara Etna** | Spanish and Greek style, green | F454 | 16S rRNA and rRNA gene | Effect of type of processing on bacterial microbiota | (Cocolin et al., 2013a) |

ITS internal transcribed spacer

Thereby, in this work our objective was to use the FoodMicrobionet database (Parente et al., 2016, 2022; Parente and Ricciardi, 2024) to provide quantitative insights on the structure of bacterial and fungal microbial communities of table olives and to identify core genera as a function of table olive elaboration. Even if an obvious shortcoming of our approach is the heterogeneity of data available (in terms of olive varieties, sampling strategies and experimental designs, gene targets and primers, sequencing platforms), we feel that the use of a standardized pipeline for raw sequence processing and an enhanced metadata structure will in part mitigate this problem, and provide transparent and reusable data and software to the scientific community. This in turn may allow a better understanding of the ecology of microorganisms which are important for the quality (including both beneficial and spoilage microorganisms) and safety of table olives and promote the development of novel microbiome-based starter cultures (Ferrocino et al., 2022; Gänzle et al., 2023).

## 2. Materials and methods

### 2.1. Database analysis

FoodMicrobionet v 5.01 (Parente and Ricciardi, 2024) includes 16 different metataxonomic studies (with a total of 833 samples) on table olives (Table 2) whose raw sequences are available from NCBI Short Read Archive (SRA). The sequences were processed with the scripts available in FoodMicrobionet repository (https://github.com/ep142/FoodMicrobionet) using computational approaches described elsewhere (Parente et al., 2022; Parente and Ricciardi, 2024).

**Table 2.**
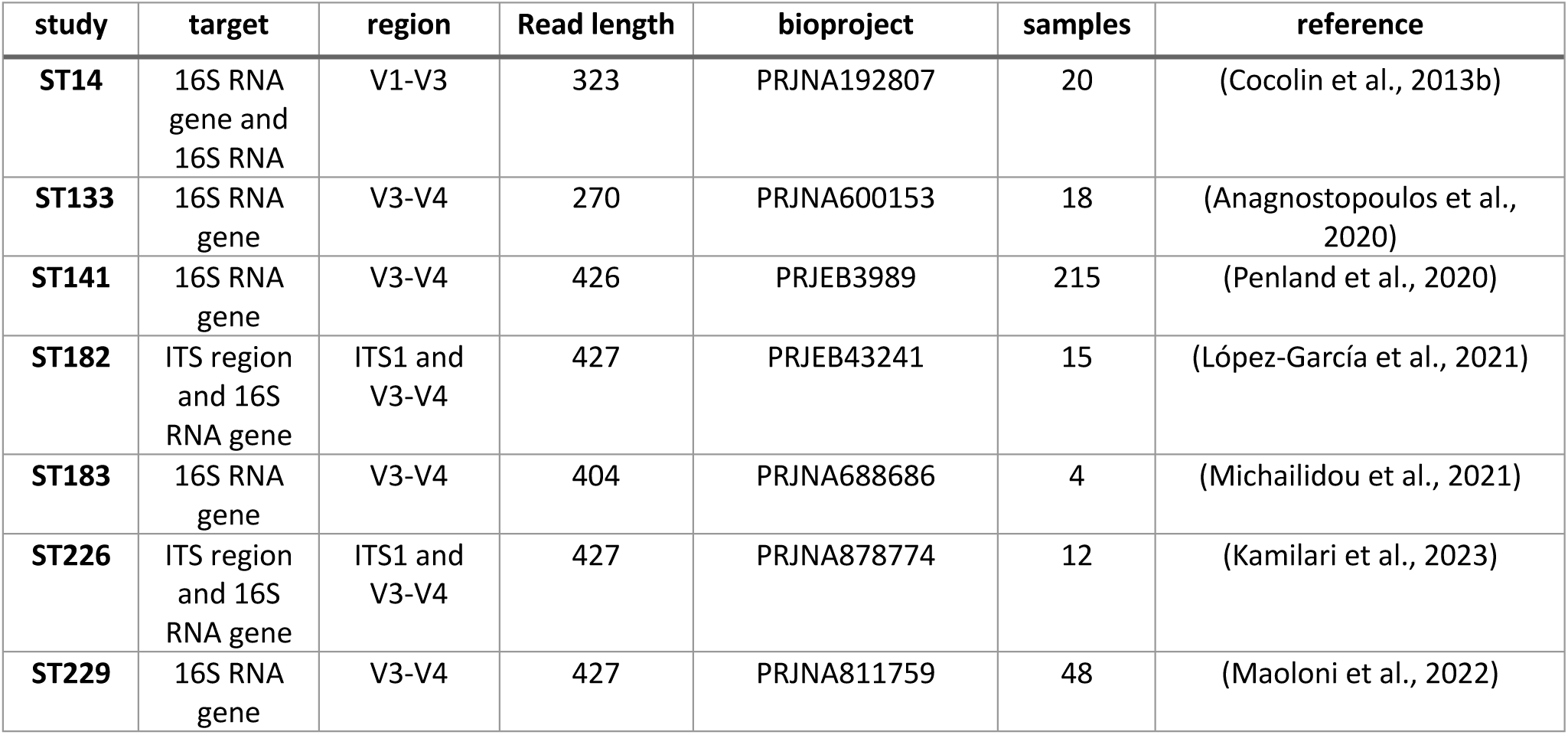
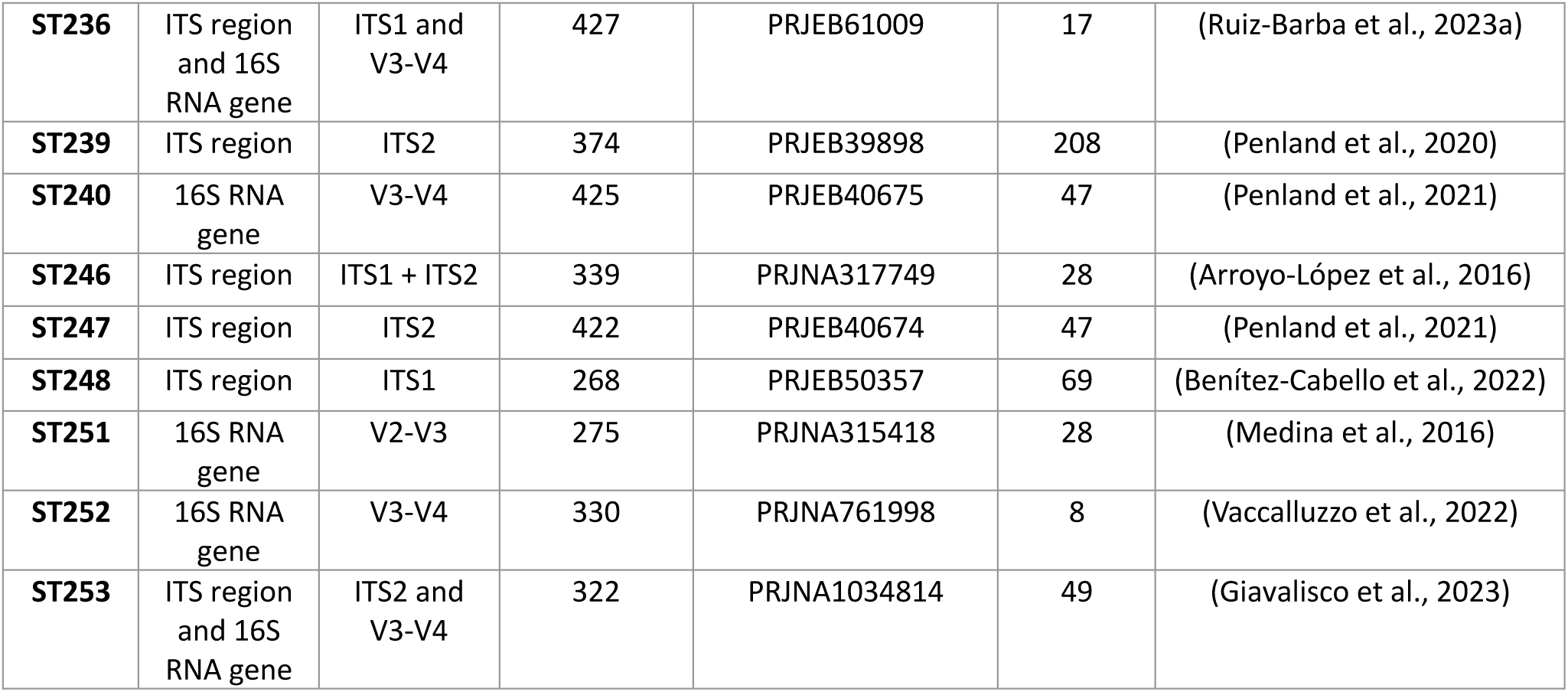
List of studies on metataxonomic table olives included in FoodMicrobionet 5.0.1 (Parente and Ricciardi, 2024) which were used in the present work.

ST253 is an unpublished study from our group, dealing with the effect of starter culture (*Lactiplantibacillus pentosus* O17) and timing of salt addition to the brine on the manufacture of black Itrana olives (Giavalisco et al., 2023). Sequences were deposited in NCBI SRA with accession number PRJNA1034814.

All studies were extracted from FoodMicrobionet using the extract_studies.R script available in the Metaolive repository (https://github.com/ep142/Metaolive) and saved as phyloseq objects (McMurdie and Holmes, 2013).

Until recently metadata models were not enforced for the deposit of sequence data in NCBI Short Read Archive (https://www.ncbi.nlm.nih.gov/biosample/docs/packages/): the metadata for studies deposited in SRA are therefore often incomplete and inconsistent and do not follow a common scheme and those available in FoodMicrobionet are not specific for table olives. We therefore assembled a complete set of metadata attributes (olive cultivar, olive variety, olive trade preparation, style, spoilage, starter addition, etc.), by using information available directly from metadata deposited in SRA or inferred from the papers describing the study, were added to the sample tables. The enriched metadata are available in our GitHub repository (https://github.com/ep142/Metaolive/tree/main/olive_FMBN).

Graphical and statistical analysis of phyloseq objects was carried out using own R 4.3.2 (R core team, 2024) scripts, available in the GitHub repository. Taxonomic re-assignment was carried out using the assignTaxonomy() and addSpecies() functions of the dada2 package (Callahan et al., 2016), using Silva 138.1 (Quast et al., 2013) and UNITE (general release 04/04/2024) (Abarenkov et al., 2023) taxonomic reference databases. Statistical and graphical treatment of the results included:

1. sample filtering (by removing samples with < 1000 sequences)
2. primary taxa filtering: sequences to mitochondria, and chloroplasts were removed from bacterial datasets; sequences assigned to fungi were removed from bacterial datasets, and vice versa;
3. taxonomic agglomeration was performed at the genus level;
4. application of a very lenient prevalence (threshold for removal < 0.01) and abundance (threshold for removal < 0.005) filter (taxa were retained if they passed both thresholds) to avoid loss of rare taxa;
5. calculation of alpha diversity
6. production of a variety of boxplots and bar plots for alpha diversity
7. ordination using non-monotonic multidimensional scaling of the Bray-Curtis distance calculated on taxa abundance data

To this end, functions from packages phyloseq, vegan (Oksanen et al., 2022) and tidyverse (Wickham et al., 2019) were used.

Detailed metadata are available in our GitHub repository (https://github.com/ep142/Metaolive).

Of the 16 studies available, 8 included data for bacteria only, 4 for fungi only, and 4 for both bacteria and fungi. However, due to inconsistencies in the deposit of sequences in SRA (in several cases the same sample was deposited with two separate biosample accessions and/or data for bacteria and fungi were deposited with different bioproject or study accessions), the same samples might be present in two studies, one for bacteria and one for fungi (by convention in FoodMicrobionet a study must have a unique bioproject accession and a sample a unique biosample accession). The number of samples per study varied from 4 to 251, with a median value of 28.

## 3. Results and Discussion

### 3.1 The bacterial microbiota of table olives

Data on bacterial communities of table olives and their production environments were obtained from 12 metataxonomic studies, whose number of samples was highly unbalanced (Supplementary Table 1). The most common target and platform are the V3-V4 region of the 16S RNA gene and Illumina, respectively; data on 16S RNA were found in one study but were removed from the analysis.

Samples were available for fruits and/or brine, or contact surface and raw materials (Supplementary Table 2 and 3). After removing samples with less than 1,000 sequences, a total of 411 samples were left for analysis (Supplementary Table 3).

The largest number of samples was available for Nyons (Penland et al., 2020, 2021) and Itrana naturally fermented black olives (Giavalisco et al., 2023).

Overall, the data we assembled from SRA cover a reasonably large number of olive varieties (12, see Supplementary Table 3) from 5 countries (France, Spain, Italy, Greece, and Cyprus; Supplementary Figure 1), including alkali treated and natural olives making this the most comprehensive study on the microbiota of table olives.

Alpha diversity indices were calculated before taxa filtering. Chao1 ranged from 2 to 100. The variability of Chao1 as a function of olive variety, stage of fermentation (0 environmental samples, 1 raw material, 2 samples during fermentation, samples at the end of fermentation), ripeness, trade preparation, sample type is shown in Figure 1.

**Figure 1.**
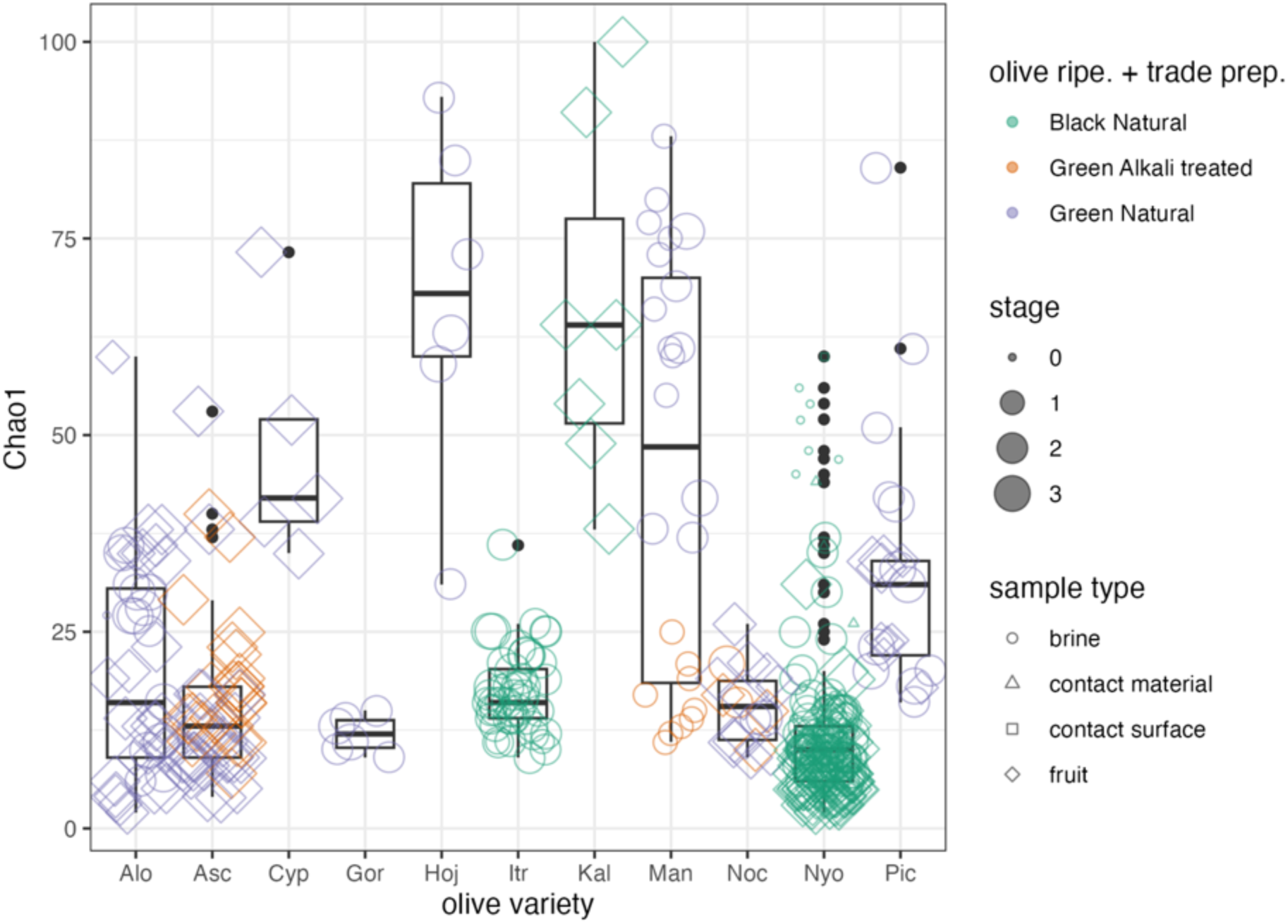
Distribution of Chao1 for bacterial communities from studies on table olives (Table 2) by olive variety, fermentation stage (1 raw materials, 2 samples during fermentation, 3 samples at the end of fermentation), olive ripeness and trade preparation, sample type.

Due to the differences in methods and experimental designs, only very general conclusions can be drawn:

- there is no evidence of a relationship between number of sequences per sample and Chao1.
- olive variety, degree of ripening and process are almost always confounded, therefore comparisons need to be done within the same variety.
- for the few studies for which the same olive variety was processed according to different trade preparations (usually Green Natural and Green Spanish style) the diversity is not consistently higher for either type.
- for the few studies for which inoculated and inoculated olives were compared, we found no significant (Kolmogorov-Smirnov two sample test) evidence of lower diversity in inoculated fermentations; median Chao1 was even higher in some cases for inoculated fermentations (see Supplementary Figure 2).
- there is no clear evidence of fermentation stage affecting the diversity (see Figure 1), i.e. diversity does not apparently decrease with increasing fermentation time due to the selection of a less diverse community adapted, for example, to high salt / high polyphenol / low nutrient conditions, although in some environmental samples (see Nyons olives) diversity is much higher than in olives or brines. The reasons for this are not clear. One reason is that, although one study (Cocolin et al., 2013a) included both 16S RNA (which may depict the active fraction within the community) and the 16S RNA gene (which may depict the structure of both the active and inactive parts of the fermentation), the number of samples was too low to make any conclusion, it is impossible to decide which portion of the microbial community as determined by amplicon targeted sequencing was active. This problem is common to many other fermentations and there is no consensus on the best approach to be used (Emerson et al., 2017; Erkus et al., 2013, 2016; Yap et al., 2021) to solve it.

After this stage taxonomic filtering and agglomeration was applied. As a result, the number of taxa was reduced from 929 (pre-taxonomic agglomeration) to 157, although the proportion of sequences retained was 99.5%. This reveals that the diversity of bacterial genera in table olives is much larger than what can be hypothesised from a single study.

A prevalence and abundance plot is shown in Supplementary Figure 3. Bacterial genera in table olives and their production environments belonged to 24 phyla, but the majority belonged to *Bacillota, Pseudomonadota, Actinobacteriota*, and *Bacteroidota*. The proportion of sequences removed by filtering varied significantly among samples: this was mostly due to the poor quality of taxonomic assignment for some samples (data not shown). However, 75% of the samples retained more than 86% of the original sequences after filtering.

We realize that the very unequal sample number per study may have significantly affected the results of filtering and the calculation of average prevalence and abundance: the majority of samples were in fact Nyons olives and their production environments (Penland et al., 2020, 2021), and Itrana olives (Giavalisco et al., 2023) (see Table 2). Given the nature of our composite dataset, this is inevitable, and subsampling to an even number of samples would greatly reduce the size of the data set and hide part of the observed diversity. Since this analysis is, in its nature, repeatable, this problem will be mitigated as the number and variety of samples for table olives in FoodMicrobionet increases. In fact, using the scripts provided in the METAolive GitHub repository (https://github.com/ep142/Metaolive), adding new data as they become available is straightforward.

Prevalence and abundance for different genera varied within different combinations of olive ripeness and trade preparation (data not shown), but a set of 31 bacterial genera whose relative abundance exceeded 0.01 in at least one sample and whose prevalence was >0.1 in at least one olive group (Green Alkali treated, Green Natural, and Black Natural) may be representative of the dominating core microbiota in table olives. These data are summarized in Figure 2, and shown in detail in Supplementary Table 4 where minimum and mean relative abundance are also shown.

**Figure 2.**
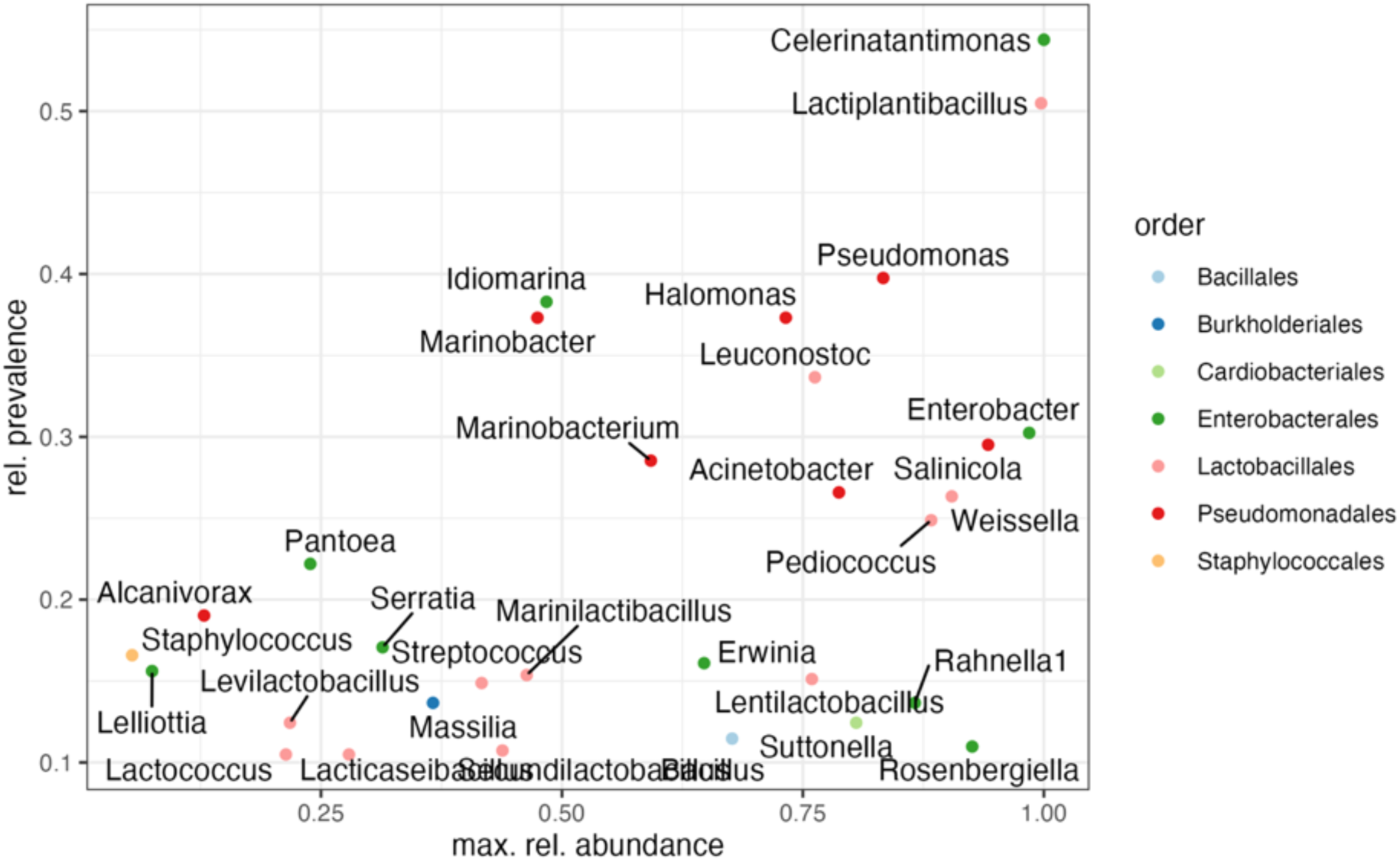
Relative prevalence and maximum relative abundance of the dominating bacterial genera in the Olive FMBN data set.

Core bacterial genera belong to two main groups: *Lactobacillales*, including Halophilic and Alkalophilic Lactic Acid Bacteria (HALAB), and Gram-negative and Gram-positive halophiles (*Enterobacterales*, *Pseudomonadales*, *Burkholderiales*, *Staphylococcales*).

Their distribution varied significantly in different olive varieties, among olives, brines and contact surfaces or materials, and at different production stages (raw materials, olives during fermentation, products at the end of fermentation). A bar plot showing the average composition of bacterial communities for olives and brines of different varieties and different trade preparations and at different stages of production is shown in Supplementary Figure 4. The 31 most abundant and dominant genera covered more than 50% of the bacterial communities for most, but not all varieties. Notable exceptions were Kalamata, Manzanilla, Aloreña, Nocellara Etnea, Hojiblanca, and Picual in which some microorganisms which were not prevalent in other varieties occurred at relatively high abundance. In Kalamata olives from Cyprus a large number of sequences was assigned to species of the genera *Streptococcus* and *Lactobacillus* (Kamilari et al., 2023), which are more frequently found in dairy products rather than in table olives. In Kalamata olives from Greece stored under modified atmosphere (Michailidou et al., 2021) a high proportion of Alpha- and Gammaproteobacteria was found at time 0, followed by an increase in the proportion of *Lentilactobacillus parafarraginis,* which was the most abundant species from 6 months onward. Further studies on this widespread variety, whose sequences were not available at the time FoodMicrobionet 5.0 was assembled, have shown that the composition of the microbiota may differ between brines and olives, and among products from different geographic areas firms (Kazou et al., 2020). *Celerinatantimonas, Lactiplantibacillus, Propionibacterium, Staphylococcus, Sediminibacterium, Leuconostoc, Sphingomonas,* and *Enterobacter* were found among the most abundant core genera by Kazou et al. (2020).

In Green Spanish style Manzanilla olives, a high proportion of *Alkalibacterium* was found. This genus was found in these olives in several studies (Arroyo-López et al., 2021; Benítez-Cabello et al., 2019; Ruiz-Barba et al., 2023a). In Nocellara Etnea green alkali treated olives (Cocolin et al., 2013a) members of the genus *Kosakonia, Citrobacter,* and more occasionally *Pantoea* and *Cobetia*, were also abundant (rel. abundance >0.10). In Aloreña de Malaga green natural olives some Actinobacteria (*Cutibacterium, Modestobacter*, *Pseudokineococcus*) were occasionally found with relative abundances >0.2 (López-García et al., 2021; Medina et al., 2018). In Picual olives from Cyprus (Anagnostopoulos et al., 2020; Kamilari et al., 2023), in addition to core genera (*Lactiplantibacillus*, *Secundilactobacillus, Pediococcus* were all relatively abundant), *Sediminibacterium,* a relatively prevalent (0.13 average prevalence) genus, was abundant (>0.3 rel. abundance in some samples, compared to 0.006 in the whole data set).

In general, we found that the composition of microbial communities detected by our pipelines using raw sequences were in good agreement with those of the original studies. Differences due to pipelines and taxonomic reference databases can be expected (Bokulich et al., 2020; Pollock et al., 2018) and DADA2 has systematically performed well in benchmarking studies (Callahan et al., 2016). The use of SILVA v138.2 allowed assignment of Amplicon Sequence Variants belonging to family *Lactobacillaceae* to the most recent taxonomic units within this family (Zheng et al., 2023) thus updating the taxonomic assignment available in older studies.

The lack of clear differences among different groups defined by combinations of ripeness and olive trade preparation is confirmed by the non-monotonic Multidimensional Scaling ordination of the Bray-Curtis distance matrix shown in Supplementary Figure 5. Rather, groups of samples within the same study (see samples for Nocellara Etnea, Itrana, Ascolana Tenera and Nyons olives for studies ST14, (Cocolin et al., 2013b); ST253, (Giavalisco et al., 2023); ST239-240, (Penland et al., 2020, 2021); and ST229, (Maoloni et al., 2022) are generally close among them. Non-monotonic Multidimensinal Scaling is an ordination technique which seeks to represent proximities in a multidimensional space in a space with typically 2 or 3 dimensions (Borg et al., 2018) while keeping the dissimilarities/distances in the new space as close to the original as possible. Therefore, sample points which lie close together in the MDS map have a similar bacterial community structure. The configuration shown in Supplementary Figure 5 confirms the results obtained from alpha-diversity analysis (see Supplementary Figure 4): selective factors which may be associated to olive ripeness (sugar concentration, polyphenol concentration, etc.) or trade preparation (alkali treatment followed by fermentation in Spanish style olives compared to fermentation alone in naturally fermented olives) are apparently not powerful enough to result in clearly separated microbial communities or to overcome confounding factors related to individual producers (differences in initial contamination, fermentation conditions, including salt concentration, temperature and atmosphere composition, etc.). Additionally, studies with several producers for the same variety are very rare and more data are needed to clarify this issue.

Addition of starter cultures affects the composition of the microbiota, and, as a consequence, the sample position in the ordination to a variable extent: differences are larger in some study (ST253; Itrana olives) than in others (ST229, Ascolana Tenera, (Maoloni et al., 2022); ST133, Picual, (Anagnostopoulos et al., 2020) possibly reflecting different abilities of the starters in colonizing fruit and brines and in affecting directly and indirectly other members of the microbiota. Maoloni et al. (2022) used a four-species LAB starter which only caused small changes in the composition of the microbiota, while a commercial *Lactipl. plantarum* starter did dominate the microbial community of cracked and, to a lesser extent, whole Picual olives, especially at the species level. The effect was also indirect: Anagnostopoulos et al. (2020) found that *Lactipl. plantarum* addition apparently promoted the growth of *Marinilactobacillus.* We were unable to replicate this finding but we did confirm that *Pediococcus* abundance was lower in started fermentations. In our work (Giavalisco et al., 2023) on Itrana olives, the addition of a *Lactipl. pentosus* starter clearly reduced the relative abundance of *Leuconostoc* and *Pseudomonadota* compared to unstarted fermentations, at least at later fermentation stages.

Spoilage does also often cause notable changes in the composition of the microbiota, as shown in the ordination of Supplementary Figure 6. Spoiled samples of Nocellara Etnea and Nyons olives clearly stand apart from unspoiled samples in the same varieties/studies.

The variability in the relative abundance and distribution of LAB and HALAB in table olives is shown in Figure 3. In this and in the other boxplots samples in which a given genus was absent are shown at the bottom of the graph. The significance of 0s in metataxonomic data has always been an issue: they could reflect lack of sensitivity (due to shallow sampling) or real absence (Lutz et al., 2022). Due to differences in sequencing depth in our dataset (we did not to apply any rarefaction; McMurdie and Holmes, 2014) we decided not to replace 0s with dummy values. As a consequence, the quantiles reflect the abundance of a given genus solely in the sample in which it was found, while its prevalence is shown in Supplementary Table 4.

**Figure 3.**
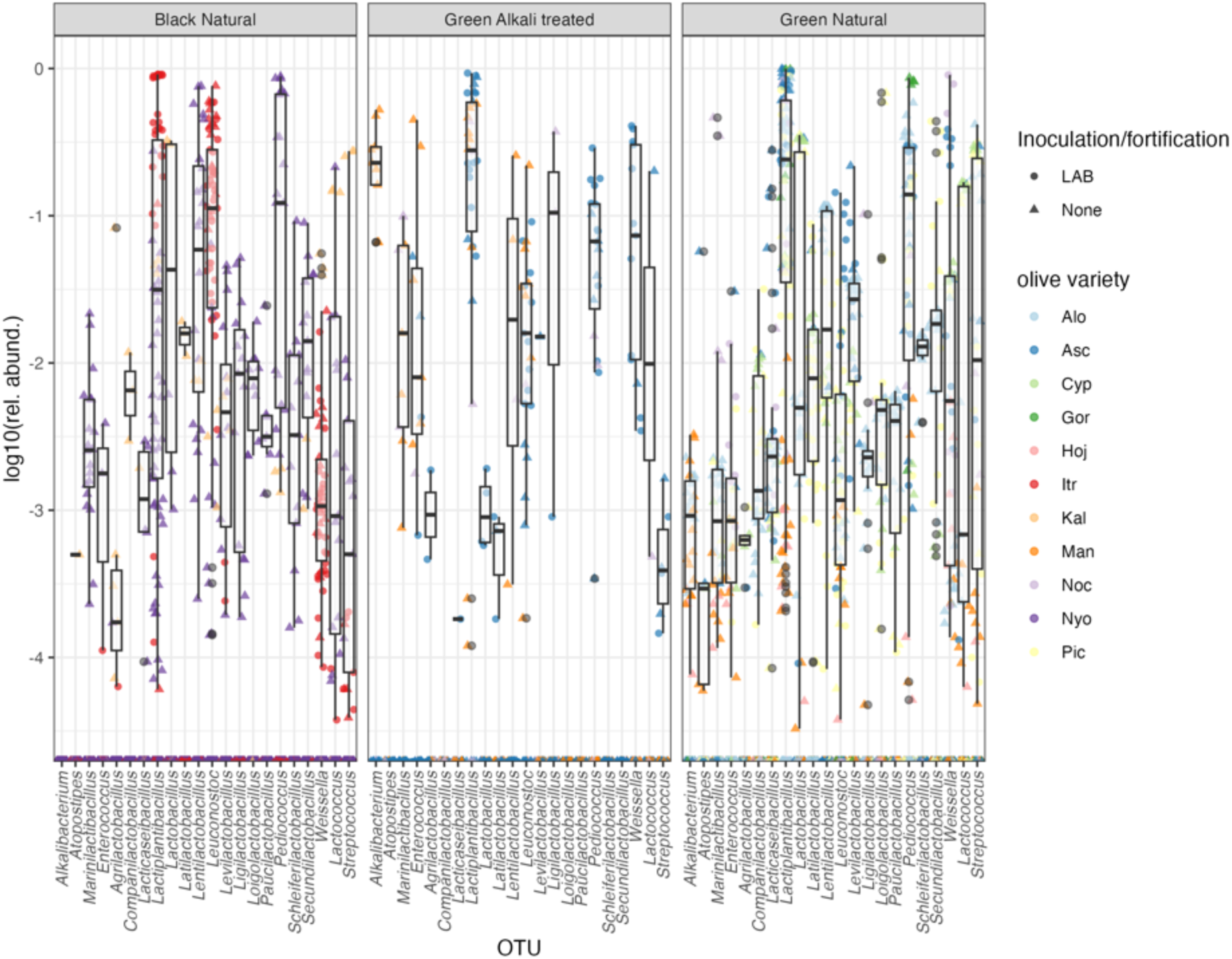
Distribution of the relative abundance of Lactic Acid Bacteria (including Halophilic and Alkalophilic LAB) in table olives. See Supplementary Table 3 for the abbreviations of table olives varieties

Homofermentative genera, with *Lactiplantibacillus* as the most prevalent and abundant genus, followed by members of the genera *Lacticaseibacillus, Pediococcus* and *Latilactobacillus,* are frequently found by amplicon targeted metagenomics in several varieties of olives (Portilha-Cunha et al., 2020; Tsoungos et al., 2023), and they are sometimes used as starters (Giavalisco et al., 2024). Other minor homofermentative genera (*Schleiferilactobacillus*, *Loigolactobacillus*) are far less prevalent but can sometimes be subdominant. Some heterofermentative genera (*Leuconostoc*, *Lentilactobacillus, Levilactobacillus, Ligilactobacillus, Secundilactobacillus, Weissella)* frequently occur at relative abundances >0.01 and can occasionally be abundant in table olives: their role is unclear (Portilha-Cunha et al., 2020; Tsoungos et al., 2023), and *Lentilactobacillus* can be involved in spoilage *(Lent. parafarraginis* and *Lent. buchneri* have been found in brines of spoiled Nyons olives; Penland et al., 2021) or produce biogenic amines. Enterococci, streptococci and lactococci are occasionally found as abundant members of microbiota in some table olive samples, but it is unclear if some of them are contaminating sequences. While *Lactococcus lactis* is in fact commonly isolated from vegetable sources (Cavanagh et al., 2015), *Streptococcus thermophilus* has low salt tolerance and is pre-eminently associated to dairy products (Iyer et al., 2010). Other genera (homofermentative *Atopostipes*, *Companilactobacillus*; heterofermentative *Paucilactobacillus*) are usually below 0.01.

HALAB (*Alkalibacterium*, *Marinilactobacillus*, but also *Halolactobacillus*) are characteristically abundant in green, alkali treated olives (Lucena-Padrós and Ruiz-Barba, 2019, 2016; Portilha-Cunha et al., 2020). *Alkalibacterium* have been found to be abundant in spoiled olive samples (Arroyo-López et al., 2021)

The distribution of the abundance of other halophilic genera is shown in Figure 4.

**Figure 4.**
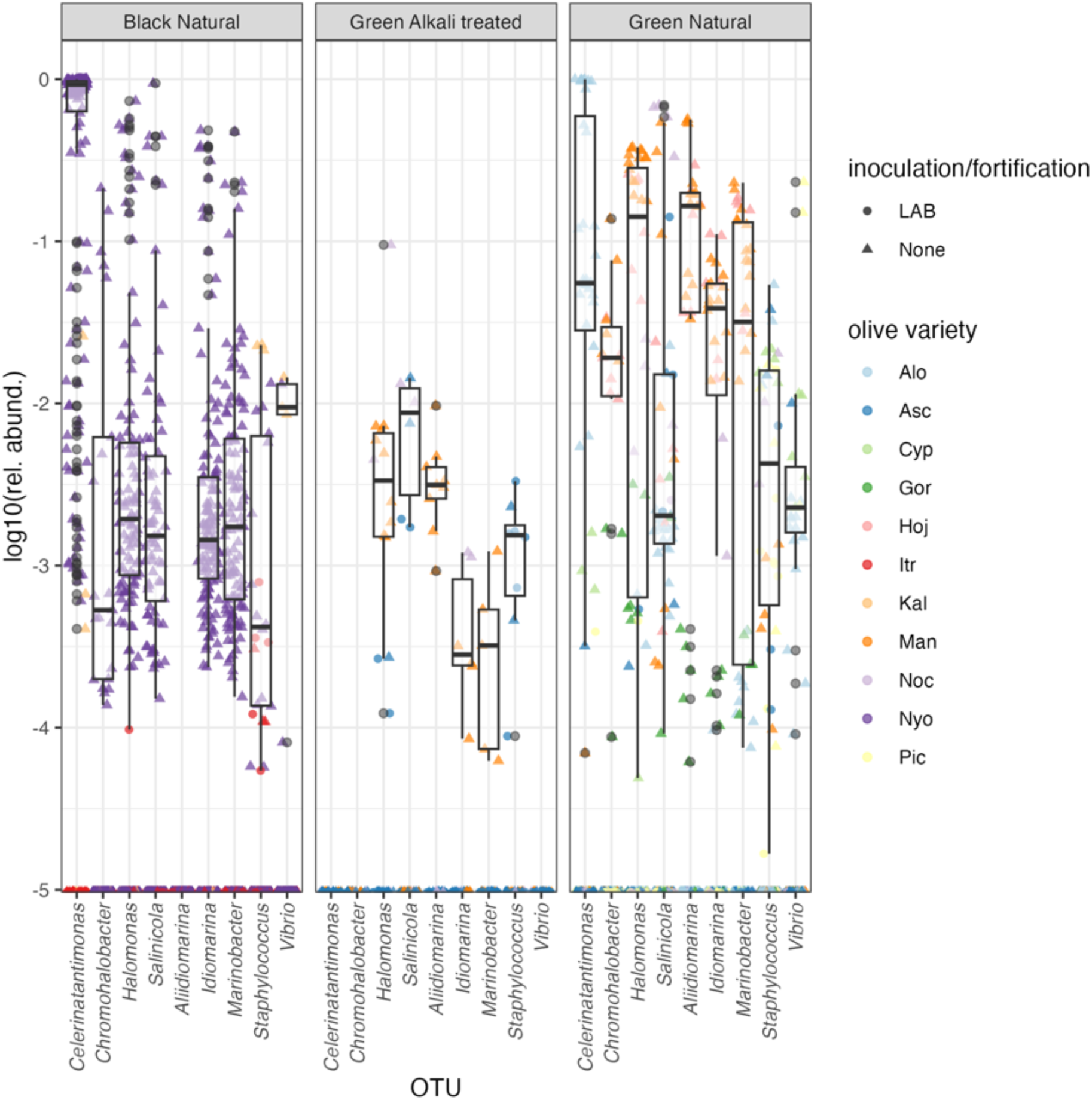
Distribution of the relative abundance of halophilic genera (other than HALAB) in table olives.

A number of halophilic *Pseudomonadota* are found in table olives, sometimes with high abundance and prevalence. The most notable genus is *Celerinatantimonas* which has been found to be abundant in Nyons black olives (Penland et al., 2021, 2020) where they are not associated with spoilage, and is also often found at relatively high abundances in green naturally fermented olives (Benítez-Cabello et al., 2020b; López-García et al., 2021; Medina et al., 2016; Rodríguez-Gómez et al., 2017). *Celerinatantimonas diazotrophica* has been specifically associated with the spoilage of Manzanilla and Gordal olives (de Castro et al., 2022; Ruiz-Barba et al., 2022). Most of the other species are associated with salt and brines. *Vibrio* have been isolated from table olives (Lucena-Padrós et al., 2015) but several *Vibrio* species are apparently unable to grow in olive fermentation brines, possibly because of inhibition by phenolic compounds (Posada-Izquierdo et al., 2021). However, they are found among the dominant members of the microbiota of Hojiblanca green Spanish style olives and might be favored by alkaline conditions with high salt; their presence in brine is not desirable (Correa-Galeote et al., 2022).

Other halophilic *Pseudomonadota* (*Alidiiomarina*, *Halomonas*, *Idiomarina*, *Marinomonas*, and *Salinicola*) are also abundant in green natural olives and in some black natural olives (Nyons olives), but less so in green alkali treated olives (Cocolin et al., 2013a; Randazzo et al., 2017; Ruiz-Barba et al., 2023a, 2023b). Fresh brines are a likely source of some of these species (Penland et al., 2021).

Because of low pH, relatively high NaCl and presence of polyphenols, table olives are a safe food (with the possible exception of olives darkened by oxidation, which need to be submitted to a botulinum cook (Campus et al., 2018)) and, to our knowledge, they have never been implicated in a food borne outbreak. Short reads used in amplicon targeted metagenomics seldom allow the resolution of closely related species and, in our study, we choose to agglomerate data at the genus level. A boxplot showing the distribution of the abundance of some genera which include human pathogens is shown in Supplementary Figure 7. Sequences assigned to *Brucella* and *Campylobacter* were only occasionally found and only at very low levels (relative abundance <0.001). On the other hand, sequences assigned to the genera *Salmonella* and *Yersinia* were found in Itrana, Ascolana Tenera, Aloreña and Hojiblanca, sometimes with relative abundances >0.01. *Listeria* was never found. The significance of these findings is unclear. First of all, there is no way of ascertaining if these sequences come from live microorganisms. Secondly, almost none of the studies in our dataset included information which could have allowed the removal of contaminating sequences (blanks, DNA measurements, mock communities) using, for example, package decontam (Davis et al., 2018). The samples of Ascolana tenera were produced with addition of herbs and spices (Maoloni et al., 2022), which may be a source of undesirable microorganisms, but the potential source of potential pathogens in other samples is unclear.

### 3.2 The fungal microbiota of table olives

Data on fungal communities for table olives are less available in public sequence repositories compared to those on bacteria. This is rather surprising, considering the importance of yeast in table olive quality and spoilage (Arroyo-López et al., 2012; Benítez-Cabello et al., 2020a; Ciafardini and Zullo, 2019; Tsoungos et al., 2023; Tufariello et al., 2019). They are available in FoodMicrobionet for 8 studies (Supplementary Table 5) from 6 countries (Cyprus, France, Greece, Italy, Perú, and Spain; Supplementary Figure 8) with a total of 453 samples (including fruits, brine, fruit+brine, raw materials and contact surfaces). Although in most cases samples were collected before or during the fermentation, in a single study packaged commercial samples from a large variety of sources was used (Benítez-Cabello et al., 2020b).

The number of samples per study was highly unbalanced (12-208, median 37.5). As for bacterial communities, most were contributed by two studies, one on black natural Nyons olives (Penland et al., 2020, 2021) and one on Itrana olives (Giavalisco et al., 2023). Samples were available for fruits and/or brine, or contact surface and raw materials. After removing samples with less than 1,000 sequences, 390 samples were left for analysis (Supplementary Table 6), which makes this the largest and more diverse combined dataset on the fungal microbiota of olives.

As for bacterial microbiota, alpha diversity indices were calculated before taxa filtering. Chao1 ranged from 2 to 100. The variability of Chao1 as a function of olive variety, stage of fermentation, ripeness, trade preparation, sample type is shown in Figure 5.

**Figure 5.**
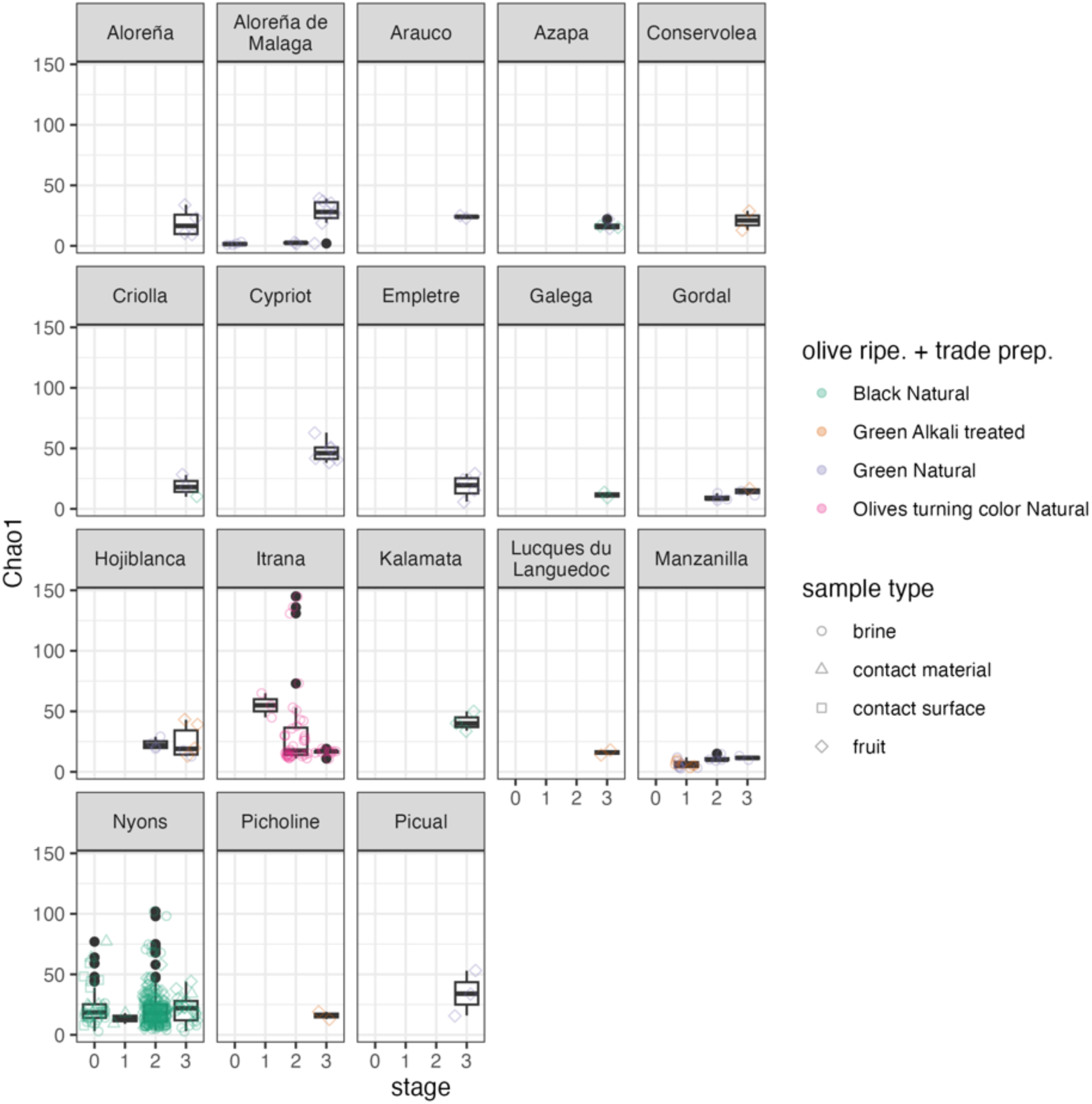
Distribution of Chao1 for fungal communities from studies on table olives (Table 2) by olive variety, fermentation stage (1 raw materials, 2 samples during fermentation, 3 samples at the end of fermentation, 0 other, including environmental samples), olive ripeness and trade preparation, sample type.

For black natural olives of the Itrana variety (Giavalisco et al., 2023) there was a clear trend for decrease of diversity from raw olives to the end of fermentation (Supplementary Figure 9), but this is almost entirely due to the higher diversity in raw, uninoculated olives and before day 25, while no significant differences were found between started and uninoculated brines (data not shown).

However, there is no evidence of a similar pattern for the few other studies for which time-course data are available (ST182, ST236, ST239), even when duration of fermentation was very long (ST239). Therefore, with data currently available we cannot conclude that fungal diversity in table olives will always decrease over time.

Taxonomic filtering and agglomeration at the genus level was performed as described for bacteria. Even if our pipeline allowed taxonomic assignment at the species level, given the differences in targets used in this study and the potential biases in taxonomic assignment related to different regions of ITS used for metataxonomic analysis (Hakimzadeh et al., 2023; Kauserud, 2023; Rué et al., 2023) the use of genus level assignment may be more prudent when comparing several studies. As a result of taxonomic filtering and agglomeration, the number of taxa was reduced from 535 to 140, and the proportion of sequences retained was 99.5%. A prevalence and abundance plot is shown in Supplementary Figure 10. Most of the genera passing the filters belonged to the phyla Ascomycota and Basidiomycota.

Prevalence and abundance for different genera varied within different combinations of olive ripeness and trade preparation (this is evident from Supplementary Figures 11-14), but a set of 31 fungi genera whose relative abundance exceeded 0.01 in at least one sample and whose prevalence was >0.1 in at least one olive group (Green Alkali treated, Green Natural, and Black natural) may be representative of the dominating core fungal microbiota in table olives (Figure 6, Supplementary Table 7).

**Figure 6.**
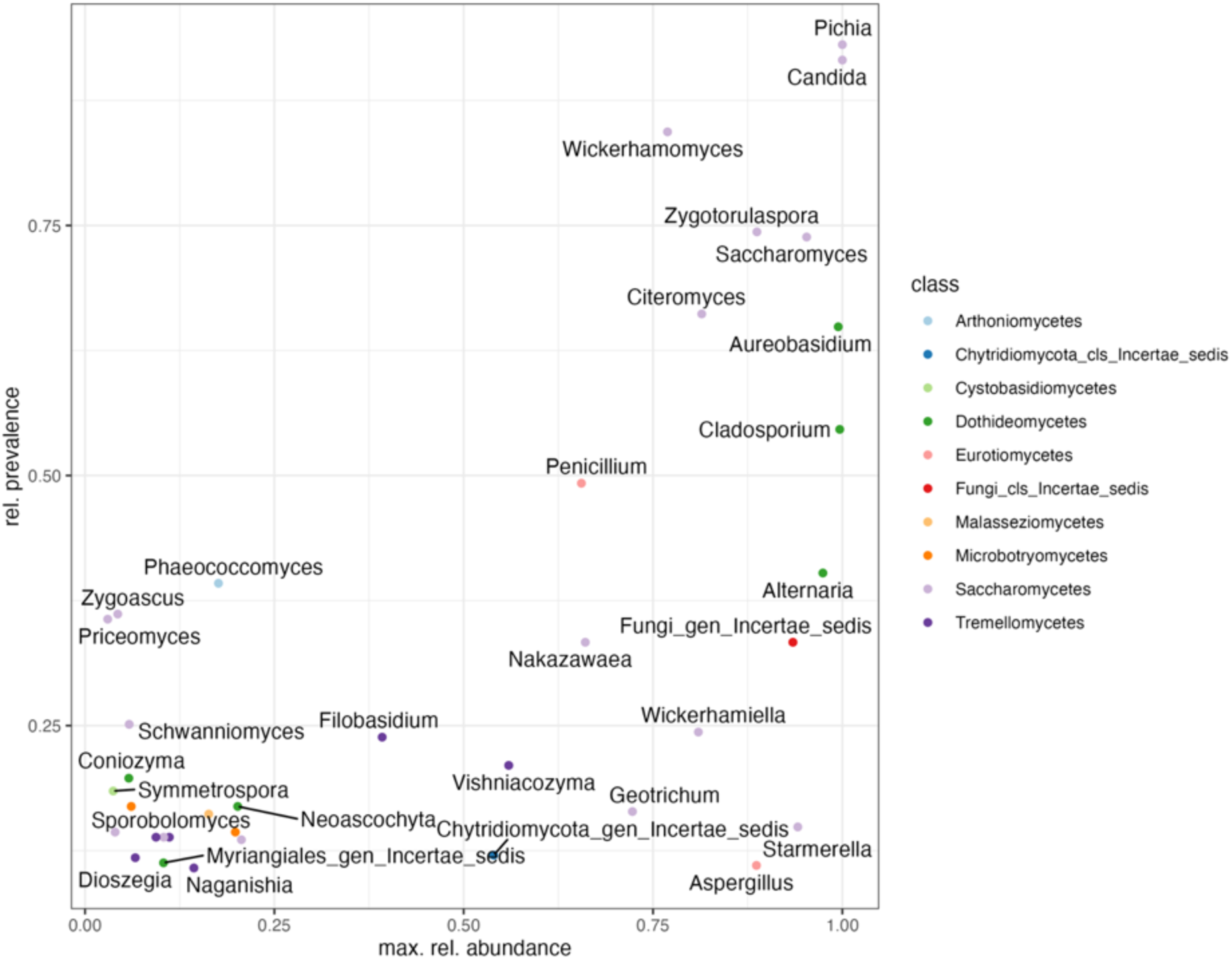
Relative prevalence and max relative abundance of the dominating fungal in the Olive FMBN data set

Given the high number of olive varieties it is impossible to show them in a single plot. The distribution of the core fungal genera is shown in Figure 7, while data on individual varieties are shown in the Supplementary Figures 11-13.

**Figure 7.**
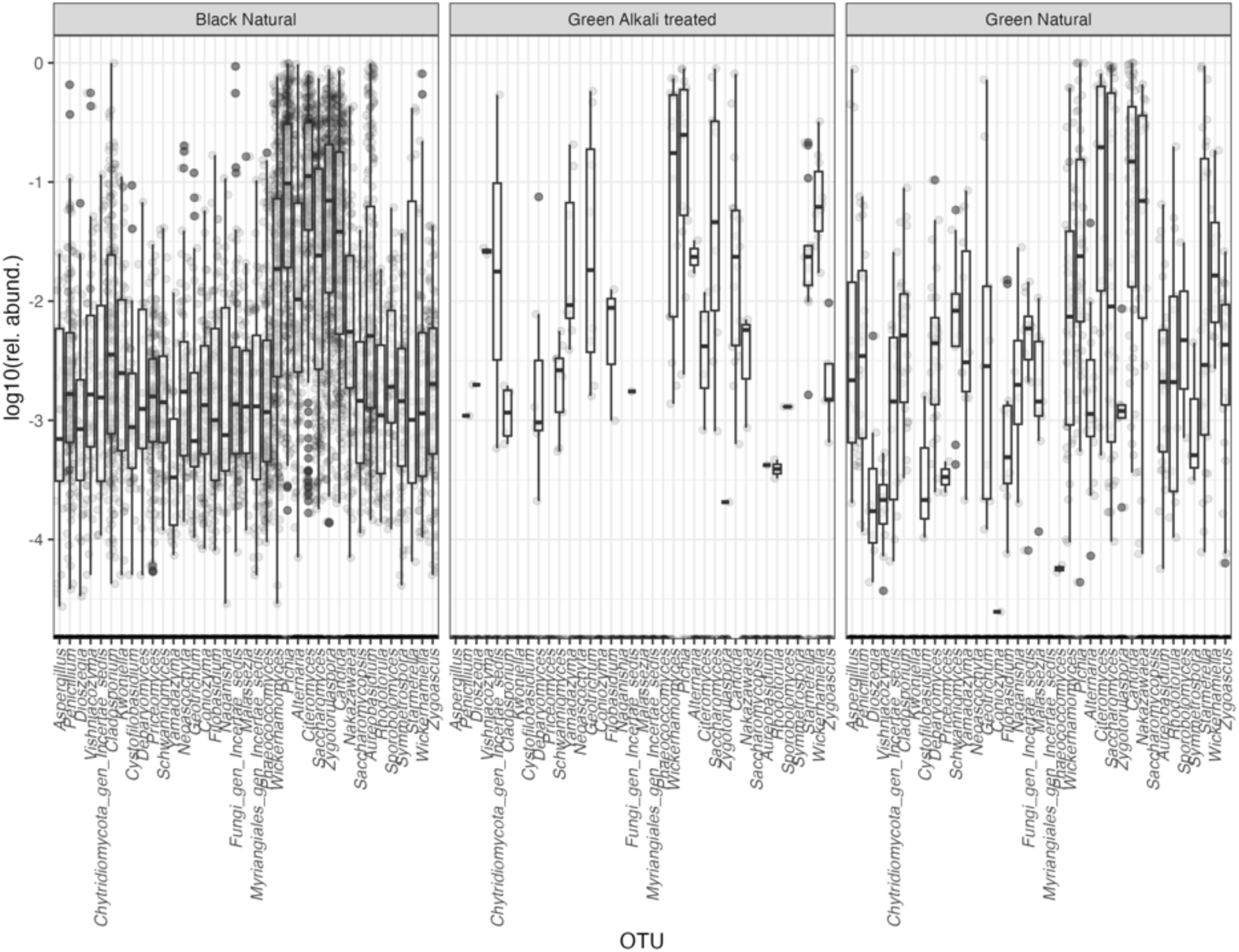
Distribution of the dominating fungal genera in the different types of table olive processing.

The distribution of abundance of fungi in table olives and their brines and food production environments was highly variable among olive groups and varieties (Supplementary Figure 14). The largest diversity was found in black natural olives, in which all core genera were found (although only *Pichia* and *Cyteromyces* had a median relative abundance >0.1). The variety of core genera found in green natural and green alkali treated olives was lower, and the most abundant genera were different (*Pichia* and *Wickerhamomyces* in green alkali treated olives, and *Pichia*, *Cyteromyces,* and *Candida* in green natural olives).

For the varieties for which samples at different time points were available (Itrana, Nyons, Hojiblanca, Manzanilla) a clear evolution of the composition of the microbiota over time is evident. The abundance of *Pichia* tended to increase over time for Nyons olives. Although this genus was also among the most abundant for the other three varieties at the end of fermentation, it was not the most abundant (*Candida* dominated in Itrana olives and *Saccharomyces* in Hojiblanca and Manzanilla).

The most abundant and prevalent fungal genera included members of the genus *Candida* (species level taxonomic assignment included *C. boidinii*, the species with the highest average relative abundance and prevalence*, C. diddensiae,* and *C. spencermartinsiae*), *Pichia* (with the two closely related species *P. membranifaciens* and *P. manshurica*) and *Wickerhamomyces* (*W. anomalus*). These species have been regularly found in culture dependent and independent studies on the fungal communities of olives, both during fermentation and in commercial packages (Anagnostopoulos and Tsaltas, 2022; Arroyo-López et al., 2012; Tsoungos et al., 2023). Members of the genus *Saccharomyces* (*S. cerevisiae*) are also prevalent and may be abundant in some varieties (Nyons, Penland et al., 2021, 2020; Itrana, Giavalisco et al., 2023). They have been frequently reported as dominating and subdominating members of the fungal microbiota of table olives (Arroyo-López et al., 2016; Kamilari et al., 2023; Kazou et al., 2020; Ruiz-Barba et al., 2023a, 2023b; Tsoungos et al., 2023; Tzamourani et al., 2022).

*Cyteromyces* (*Cyt. nyonsensis*) was found in both black and green natural olives with relatively high abundance (see below). It was first described in Nyons olives (Casaregola et al., 2013), but is also found in Taggiasca olives (Traina et al., 2024) and in black dry salted olives (Gounari et al., 2023).

Other yeast genera which have been commonly associated to table olives and olive brines (Arroyo-López et al., 2012; Tsoungos et al., 2023) like *Dekkera, Nakazawea, Zygotorulaspora, Wickerhamiella, Starmerella, Schwanniomyces*, are much less abundant and prevalent.

Other like *Dioszegia*, *Kwoniella* and *Vishniacozyma* are found occasionally and at lower abundances and may be associated to the phylloplane rather than to the fermentation (Ferluga et al., 2024; Wang et al., 2008; Zhu et al., 2021) (see also the Omnicrobe database https://omnicrobe.migale.inrae.fr/index).

Yeast-like fungi (*Aureobasidium*) and molds (*Penicillium, Cladosporium* and *Aspergillus*) may have the same origin, although they are also associated with spoilage (Arroyo-López et al., 2016; Sánchez et al., 2022).

Overall, these figures confirm that there is very little in terms of common patterns of the composition of fungal microbiota among different combinations of olive ripeness and olive trade preparation. Although some genera (*Pichia, Candida, Wickerhamomyces* and *Saccharomyces*) tend to be more abundant and prevalent than others in almost all varieties, their abundance can be highly variable among different varieties in the same group and, for the few cases in which data are available, large changes are observed between the earliest stages of fermentation (raw olives, olives at the beginning of fermentation) and intermediate stages. The latter tend to be similar to late stages. In addition, in the few cases in which data for both fruits and brine are available, differences existed between the two niches.

This is confirmed by the ordination plots shown in Supplementary Figures 15 and 16. For individual studies, a clear separation of samples can be found, and this can be related to factors for a given study. For example, evolution over time is an important factor for Itrana, Nyons, Hojiblanca and Manzanilla, see above; differences among unfermented olive fruits, contact surfaces and materialas and fermented brines or olives for Nyons; spoilage for Nyons olives.

## 4. Conclusions

In this paper, we have used in a transparent and reproducible manner, public metataxonomic data sets on the composition and dynamics of microbial communities in table olives to obtain a comprehensive view of the bacterial and fungal core microbiota of table olives.

Even if the microbiota of table olives has been thoroughly reviewed in recent years (Anagnostopoulos and Tsaltas, 2022; Campus et al., 2018; Corsetti et al., 2012; Perpetuini et al., 2020; Portilha-Cunha et al., 2020; Tsoungos et al., 2023), the quantitative insights provided in this paper significantly advance our knowledge of the main microbiota of different trade preparations of table olives.

On the other hand, the data we have collected point to some limitations of the experimental approaches used so far and future research needs.

Table olive fermentation is, in most cases, an artisanal process. This may explain the variability of the composition of the microbiota even within the same olive variety, which may be controlled, in part, by modern technologies for decontamination of the fruits, improved control of the fermentation process and starter addition (Campus et al., 2018). Part of the variability observed in our study may be due to the diversity of type of samples (brines, whole olives, homogenized olives, combinations of olives and brines), to differences in DNA extraction and purification procedures and other differences in wet laboratory approaches and sequencing platform (here, a common analysis pipeline was used for the analysis of raw sequences). It is well known that the microbiota of the surface of the olives and of the brines may differ (López-García et al., 2022) and it would be highly desirable if the scientific community working on table olives would agree on common protocols and sampling approaches.

Moreover, most of the metataxonomic studies available in the literature (Table 1) are small and narrowly focused. We feel that larger and more structured studies are needed to identify more reliably the microbiota associated with the fermentation of individual varieties and to relate its composition and dynamics to contamination and dispersal and to selective forces operating during the fermentation.

More and more diverse data would also be needed for the development of machine-learning approaches for the identification of specific subsets of the microbiota which are reliably associated to specific varieties or to distinguish between PDO and non-PDO processes, or to identify the microbial spoilage associations (Arroyo-López et al., 2021; Montaño et al., 1992; Penland et al., 2021).

Use of LAB and yeast starters does indeed improve the quality of table olives, as pointed out in several recent reviews (Anagnostopoulos and Tsaltas, 2022; Campus et al., 2018; Giavalisco et al., 2024). The starter cultures tested vary in their composition (Giavalisco et al. 2024): some are very simple (a single strain of a single species) and their development has been empirical. The rational design of complex, microbiome-based starters which mimic the composition and microbial interactions in the original microbial community requires both the knowledge of the composition and dynamics of microbial communities and the ability to track the survival and growth of individual strains during fermentation (Ferrocino et al., 2022; Gänzle et al., 2023). Our study does provide a more comprehensive view of the potentially beneficial genera which dominate different olive fermentations. On the other hand, amplicon targeted approaches have a limited ability to evaluate the impact of starter culture addition (see Supplementary Figures 5 and 15), because of limitations in taxonomic resolution. When cultivation based, strain typing approaches are used (Martorana et al., 2015) population dynamics are evident. Long read, 3^rd^ generation sequencing approaches can greatly improve the taxonomic resolution of metataxonomic studies (Srinivas et al., 2025). One recent, small study, has been published on the use of Oxford Nanopore Technology for the study of the microbiota of Kalamata black olives (Vougiouklaki et al., 2024). More data are needed to understand if the cost/benefit ratio of using this approach is favourable.

Evaluation of the microbial diversity of table olive fermentations is extremely important to understand strain level dynamics and the role of selection and drift in shaping, over time, the metagenome of table olive fermentations. Amplicon targeted approaches do not have enough resolution for this purpose. The number of shotgun metagenomic studies on food microbiota has been rapidly increasing and large databases are being created to allow the exploitation of this wealth of information (Carlino et al., 2024). To our knowledge, a single study reporting shotgun metagenomic data on table olives exists (Soto-Giron et al., 2021) and more data are clearly needed. Finally, comparative metagenomic studies on the main actors of table olive fermentations, such as *Lactipl. pentosus* (Page et al., 2023) would be extremely useful in analysing the patterns of domestication of these species and understanding the relationships, if any, between selective patterns associated with specific olive fermentation processes and the dominance of individual strains.

Another major issue is the detection of active fractions of the microbial community. To our knowledge, a single study used both rRNA and the 16S RNA gene as a target (Cocolin et al., 2013b), and concluded that the composition of the bacterial communities detected with both methods was similar, with the exception of one fruit sample. It is impossible to say if this is the case for all fermentations and it is well known in other food fermentations that the structure of active and inactive microbial communities may differ (Erkus et al., 2016, 2013).

Even with all these limitations in mind we feel that our work provides a significant contribution to the understanding of table olive microbiology. Our repository is publicly available (https://doi.org/10.5281/zenodo.13729346), as are the data and software used to generate it. Our data, software and analyses are therefore reproducible and reusable, and we feel they conform to the principles of FAIR (Findable, Accessible, Interoperable, Reproducible) paradigm of open science (Wilkinson, 2016). The availability of large and well annotated repositories is key to the development of machine-learning approaches for pattern recognition (Abdil et al., 2025; Kumar et al, 2024) and may contribute, as more data become available, to knowledge and applications in the table olives production.

## Acknowledgements.

This work was carried out within the PRIN 2022 Project METAOlive 2022NN28ZZ and received funding from Ministero dell’Università e della Ricerca (Rome) and the European Union Next-GenerationEU, CUP C53D23005460006 (PIANO NAZIONALE DI RIPRESA E RESILIENZA (PNRR) – MISSIONE 4 COMPONENTE 2, INVESTIMENTO 1.4 – D.D. 1048 14/07/2023).

