## Supplementary figures and tables for "Determining the core bacterial and fungal genera in table olive fermentations"

Supplementary tables and figures

**Supplementary Table 1** Studies including only data on bacterial communities of table olives and their production environments in FoodMicrobionet 5.0 (see main text for references).

| study | reference | target | region | samples | platform |
|---|---|---|---|---|---|
| **ST14** | Cocolin et al., 2013 | 16S RNA gene and 16S RNA | V1-V3 | 20 | 454 GS |
| **ST251** | Medina et al., 2016 | 16S RNA gene | V2-V3 | 28 | 454 GS |
| **ST133** | Anagnostopoulos et al., 2020 | 16S RNA gene | V3-V4 | 18 | Illumina |
| **ST141** | Penland et al., 2020 | 16S RNA gene | V3-V4 | 215 | Illumina |
| **ST240** | Penland et al., 2020 | 16S RNA gene | V3-V4 | 47 | Illumina |
| **ST182** | López-García et al., 2021 | ITS region and 16S RNA gene | ITS1 and V3-V4 | 15 | Illumina |
| **ST183** | Michailidou et al., 2021 | 16S RNA gene | V3-V4 | 4 | Illumina |
| **ST229** | Maoloni et al., 2022 | 16S RNA gene | V3-V4 | 48 | Illumina |
| **ST252** | Vacalluzzo et al., 2022 | 16S RNA gene | V3-V4 | 8 | Illumina |
| **ST226** | Kamilari et al., 2023 | ITS region and 16S RNA gene | ITS1 and V3-V4 | 12 | Illumina |
| **ST236** | Ruiz-Barba et al., 2023 | ITS region and 16S RNA gene | ITS1 and V3-V4 | 17 | Illumina |
| **ST253** | Giavalisco et al., 2023 | ITS region and 16S RNA gene | ITS2 and V3-V4 | 49 | Illumina |

**Supplementary Table 2.** Statistics on the type of samples available by olive trade preparation, studies with data on bacterial communities of table olives.

| Olive trade preparation | Sample type | n |
|---|---|---|
| **Alkali treated olives** | brine | 13 |
| **Alkali treated olives** | fruit | 30 |
| **Natural olives** | brine | 228 |
| **Natural olives** | contact material | 10 |
| **Natural olives** | contact surface | 6 |
| **Natural olives** | fruit | 212 |

**Supplementary Table 3.** Samples with data on bacterial microbiota, post sample filtering, by olive trade preparation, olive variety and sample type.

| Olive trade preparation | Olive cultivar | Olive variety | Olive variety abbr. | Olive ripeness | Sample type | n |
|---|---|---|---|---|---|---|
| **Alkali treated olives** | Ascolana tenera | Ascolana tenera | Asc | Green olives | fruit | 24 |
| **Alkali treated olives** | Manzanilla | Manzanilla | Man | Green olives | brine | 9 |
| **Alkali treated olives** | Nocellara Etnea | Nocellara Etnea | Noc | Green olives | brine | 4 |
| **Alkali treated olives** | Nocellara Etnea | Nocellara Etnea | Noc | Green olives | fruit | 6 |
| **Natural olives** | Aloreña | Aloreña de Malaga | Alo | Green olives | brine | 16 |
| **Natural olives** | Aloreña | Aloreña de Malaga | Alo | Green olives | fruit | 27 |
| **Natural olives** | Ascolana tenera | Ascolana tenera | Asc | Green olives | fruit | 25 |
| **Natural olives** | Cypriot | Cypriot | Cyp | Green olives | fruit | 5 |
| **Natural olives** | Gordal | Gordal | Gor | Green olives | brine | 6 |
| **Natural olives** | Hojiblanca | Hojiblanca | Hoj | Green olives | brine | 6 |
| **Natural olives** | Itrana | Itrana | Itr | Olives turning color | brine | 48 |
| **Natural olives** | Kalamata | Kalamata | Kal | Black olives | fruit | 7 |
| **Natural olives** | Manzanilla | Manzanilla | Man | Green olives | brine | 15 |
| **Natural olives** | Nocellara Etnea | Nocellara Etnea | Noc | Green olives | brine | 4 |
| **Natural olives** | Nocellara Etnea | Nocellara Etnea | Noc | Green olives | fruit | 14 |
| **Natural olives** | Picual | Picual | Pic | Green olives | brine | 12 |
| **Natural olives** | Picual | Picual | Pic | Green olives | fruit | 9 |
| **Natural olives** | Tanche | Nyons | Nyo | Black olives | brine | 121 |
| **Natural olives** | Tanche | Nyons | Nyo | Black olives | contact material | 10 |
| **Natural olives** | Tanche | Nyons | Nyo | Black olives | contact surface | 6 |
| **Natural olives** | Tanche | Nyons | Nyo | Black olives | fruit | 125 |

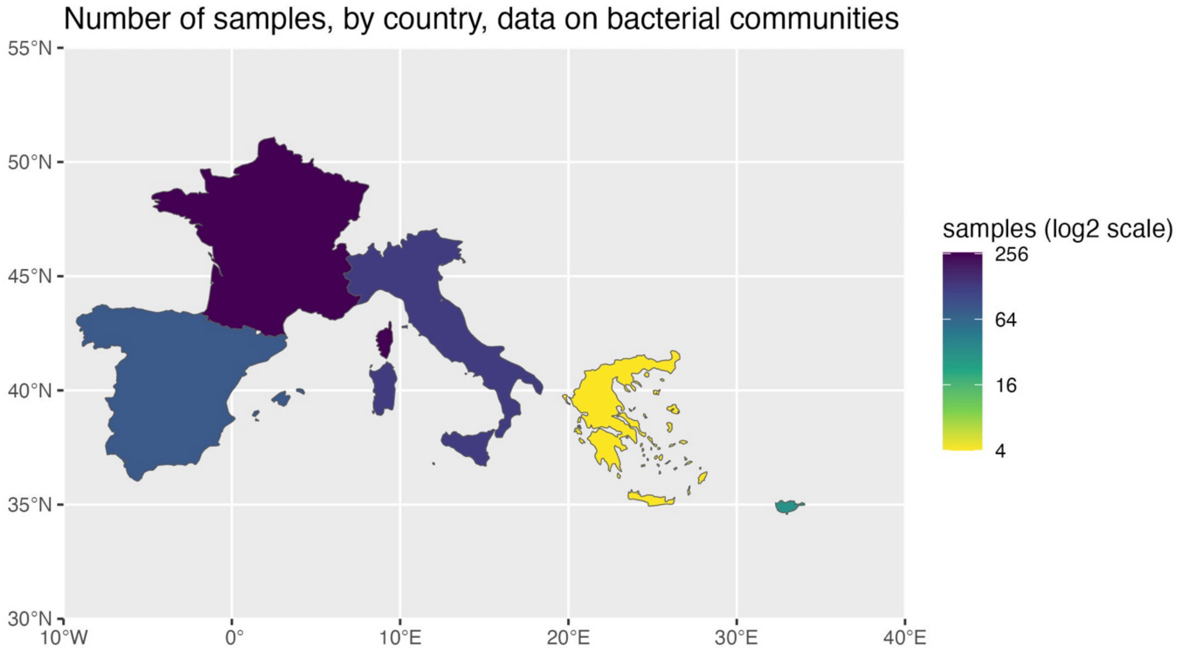


**Supplementary Figure 1**. Distribution of samples by country, data on bacterial communities of table olives.

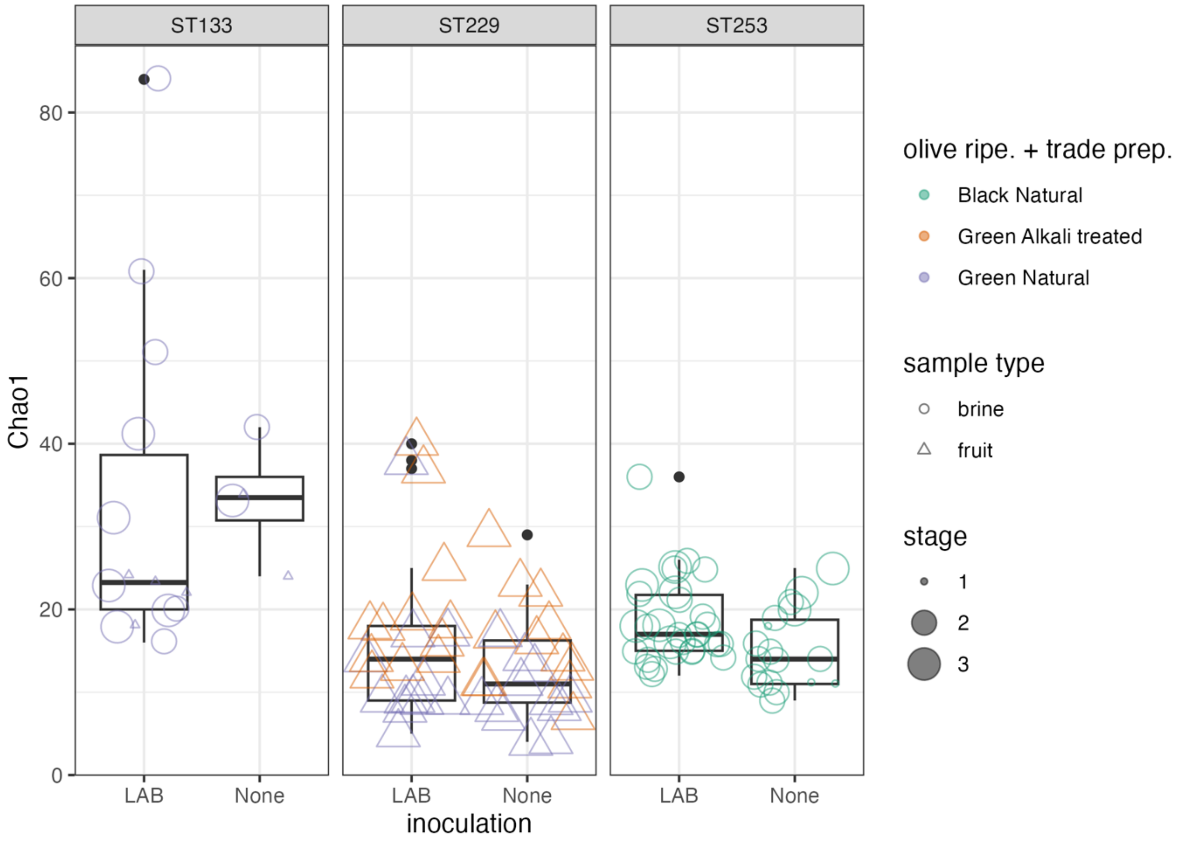


**Supplementary Figure 2.** Chao1 index for bacterial communities of table olives for studies ST133 (Picual), ST229 (Ascolana Tenera) and ST253 (Itrana).

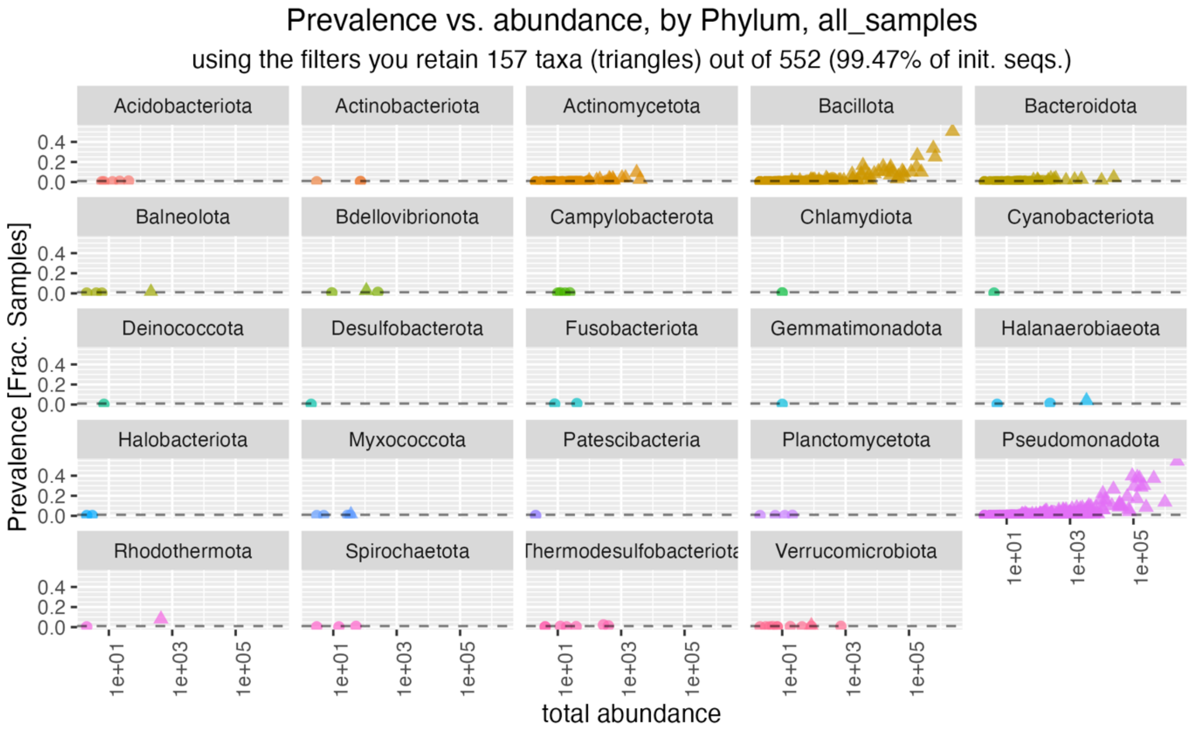


**Supplementary Figure 3.** A prevalence and abundance plot for bacterial genera in table olives, METAOlive dataset.

**Supplementary Table 4**. Prevalence and abundance data for core bacterial genera in table olives.

| Genus | Order | Prevalence | Total Abundance | Min rel. ab. | Max. rel. ab. | Mean. rel. ab. | rel. prev. |
|---|---|---|---|---|---|---|---|
| ***Celerinatantimonas*** | Enterobacterales | 223 | 2386030 | 0 | 1.00 | 2.26E-01 | 0.54 |
| ***Lactiplantibacillus*** | Lactobacillales | 207 | 2353873 | 0 | 1.00 | 2.22E-01 | 0.50 |
| ***Leuconostoc*** | Lactobacillales | 138 | 583905 | 0 | 0.76 | 5.52E-02 | 0.34 |
| ***Halomonas*** | Pseudomonadales | 153 | 436950 | 0 | 0.73 | 4.13E-02 | 0.37 |
| ***Rahnella*** | Enterobacterales | 56 | 982024 | 0 | 0.87 | 9.28E-02 | 0.14 |
| ***Pediococcus*** | Lactobacillales | 102 | 670001 | 0 | 0.88 | 6.33E-02 | 0.25 |
| ***Idiomarina*** | Enterobacterales | 157 | 131651 | 0 | 0.48 | 1.24E-02 | 0.38 |
| ***Marinobacter*** | Pseudomonadales | 153 | 155329 | 0 | 0.47 | 1.47E-02 | 0.37 |
| ***Pseudomonas*** | Pseudomonadales | 163 | 91866 | 0 | 0.83 | 8.68E-03 | 0.40 |
| ***Enterobacter*** | Enterobacterales | 124 | 210802 | 0 | 0.98 | 1.99E-02 | 0.30 |
| ***Marinobacterium*** | Pseudomonadales | 117 | 174222 | 0 | 0.59 | 1.65E-02 | 0.29 |
| ***Salinicola*** | Pseudomonadales | 121 | 105078 | 0 | 0.94 | 9.93E-03 | 0.30 |
| ***Weissella*** | Lactobacillales | 108 | 182679 | 0 | 0.90 | 1.73E-02 | 0.26 |
| ***Acinetobacter*** | Pseudomonadales | 109 | 23394 | 0 | 0.79 | 2.21E-03 | 0.27 |
| ***Pantoea*** | Enterobacterales | 91 | 10820 | 0 | 0.24 | 1.02E-03 | 0.22 |
| ***Serratia*** | Enterobacterales | 70 | 128068 | 0 | 0.31 | 1.21E-02 | 0.17 |
| ***Lentilactobacillus*** | Lactobacillales | 62 | 172116 | 0 | 0.76 | 1.63E-02 | 0.15 |
| ***Alcanivorax*** | Pseudomonadales | 78 | 63547 | 0 | 0.13 | 6.01E-03 | 0.19 |
| ***Staphylococcus*** | Staphylococcales | 68 | 3525 | 0 | 0.05 | 3.33E-04 | 0.17 |
| ***Erwinia*** | Enterobacterales | 66 | 12895 | 0 | 0.65 | 1.22E-03 | 0.16 |
| ***Lelliottia*** | Enterobacterales | 64 | 9386 | 0 | 0.07 | 8.87E-04 | 0.16 |
| ***Marinilactibacillus*** | Lactobacillales | 63 | 15254 | 0 | 0.46 | 1.44E-03 | 0.15 |
| ***Streptococcus*** | Lactobacillales | 61 | 26375 | 0 | 0.42 | 2.49E-03 | 0.15 |
| ***Massilia*** | Burkholderiales | 56 | 13255 | 0 | 0.37 | 1.25E-03 | 0.14 |
| ***Suttonella*** | Cardiobacteriales | 51 | 36634 | 0 | 0.81 | 3.46E-03 | 0.12 |
| ***Levilactobacillus*** | Lactobacillales | 51 | 25301 | 0 | 0.22 | 2.39E-03 | 0.12 |
| ***Secundilactobacillus*** | Lactobacillales | 44 | 61660 | 0 | 0.44 | 5.83E-03 | 0.11 |
| ***Rosenbergiella*** | Enterobacterales | 45 | 35423 | 0 | 0.93 | 3.35E-03 | 0.11 |
| ***Bacillus*** | Bacillales | 47 | 4129 | 0 | 0.68 | 3.90E-04 | 0.11 |
| ***Lactococcus*** | Lactobacillales | 43 | 16500 | 0 | 0.21 | 1.56E-03 | 0.10 |
| ***Lacticaseibacillus*** | Lactobacillales | 43 | 10778 | 0 | 0.28 | 1.02E-03 | 0.10 |

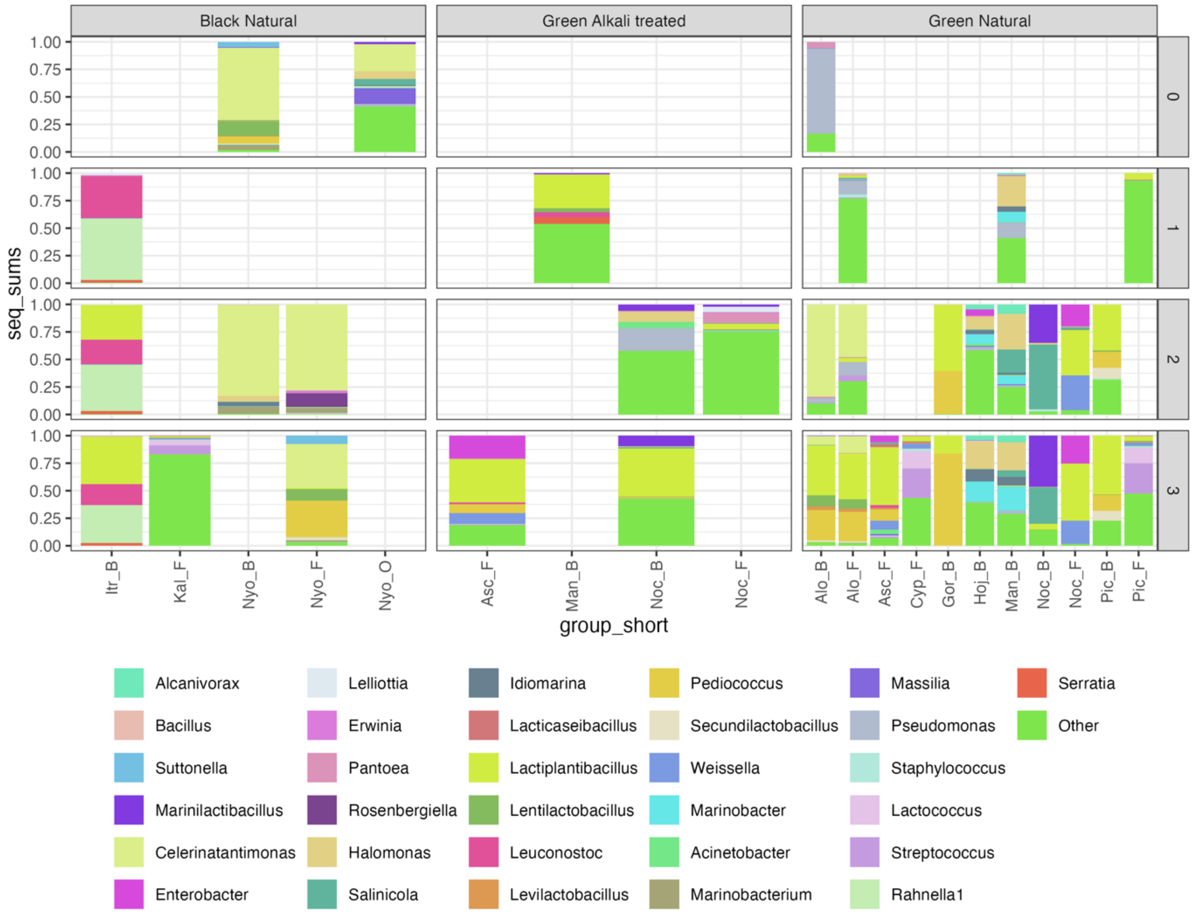


**Supplementary Figure 4**. Average composition of bacterial communities of different varieties of table olives and fermentation brine belonging to the METAOlive FMBN data set. The core genera are shown; the others are pooled under the group “Other”. Samples are divided by combination of olive ripeness and trade preparation and fermentation stage (1 beginning of fermentation, 2 samples during fermentation, 3 samples at the end of fermentation, 0 other, including environmental samples). B = brine, F = fruit. See Supplementary Table 3 for abbreviations of olive varieties.

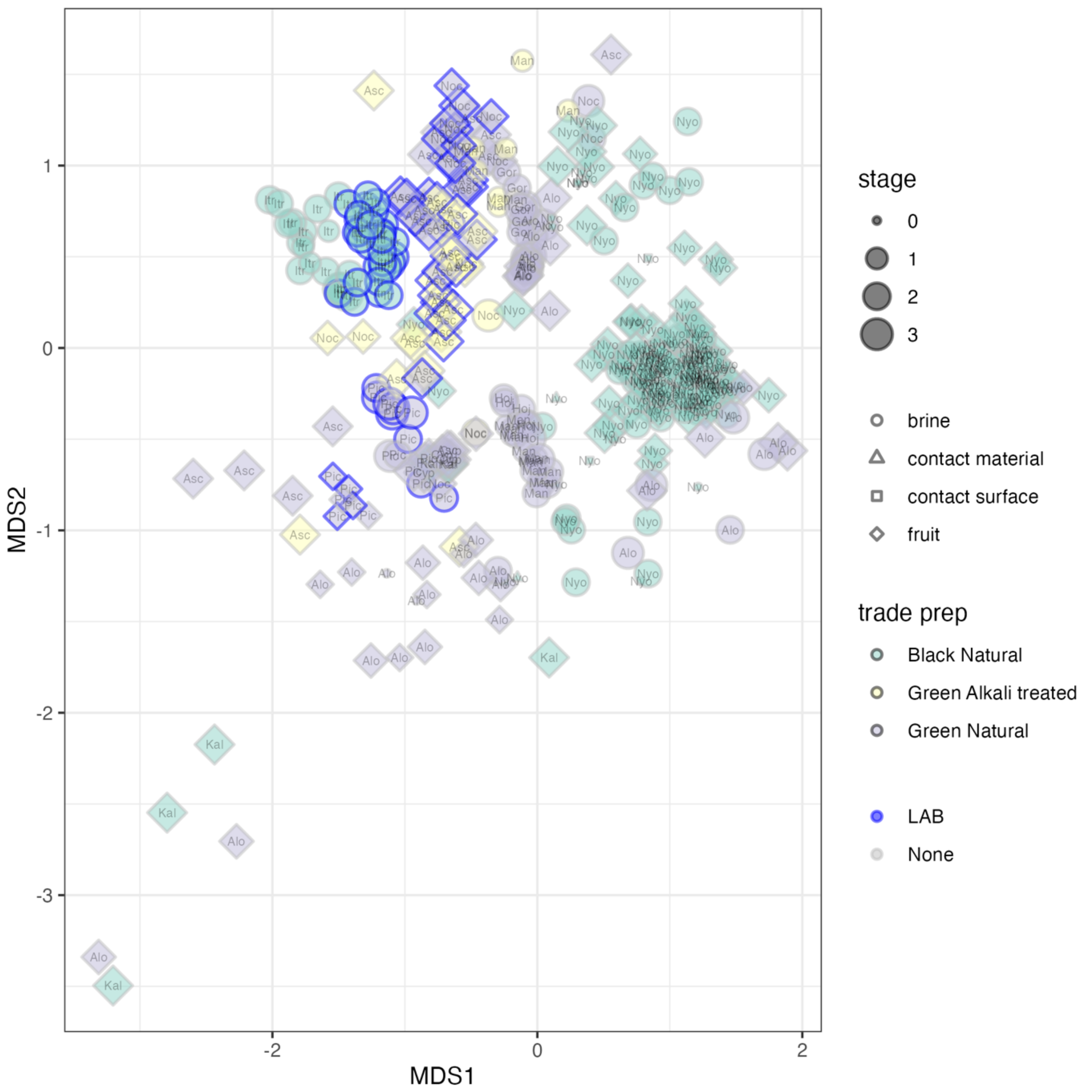


**Supplementary Figure 5**. Ordination showing the sample coordinates after non-monotonic Multidimensional Scaling of the Bray-Curtis distance matrix for the composition of bacterial microbiota of samples belonging to the METAOlive FMBN data set after prevalence and abundance filtering. Samples are divided by combination of olive ripeness and trade preparation and fermentation stage (1 beginning of fermentation, 2 samples during fermentation, 3 samples at the end of fermentation, 0 other, including environmental samples). B = brine, F = fruit. See Supplementary Table 3 for abbreviations of olive varieties.

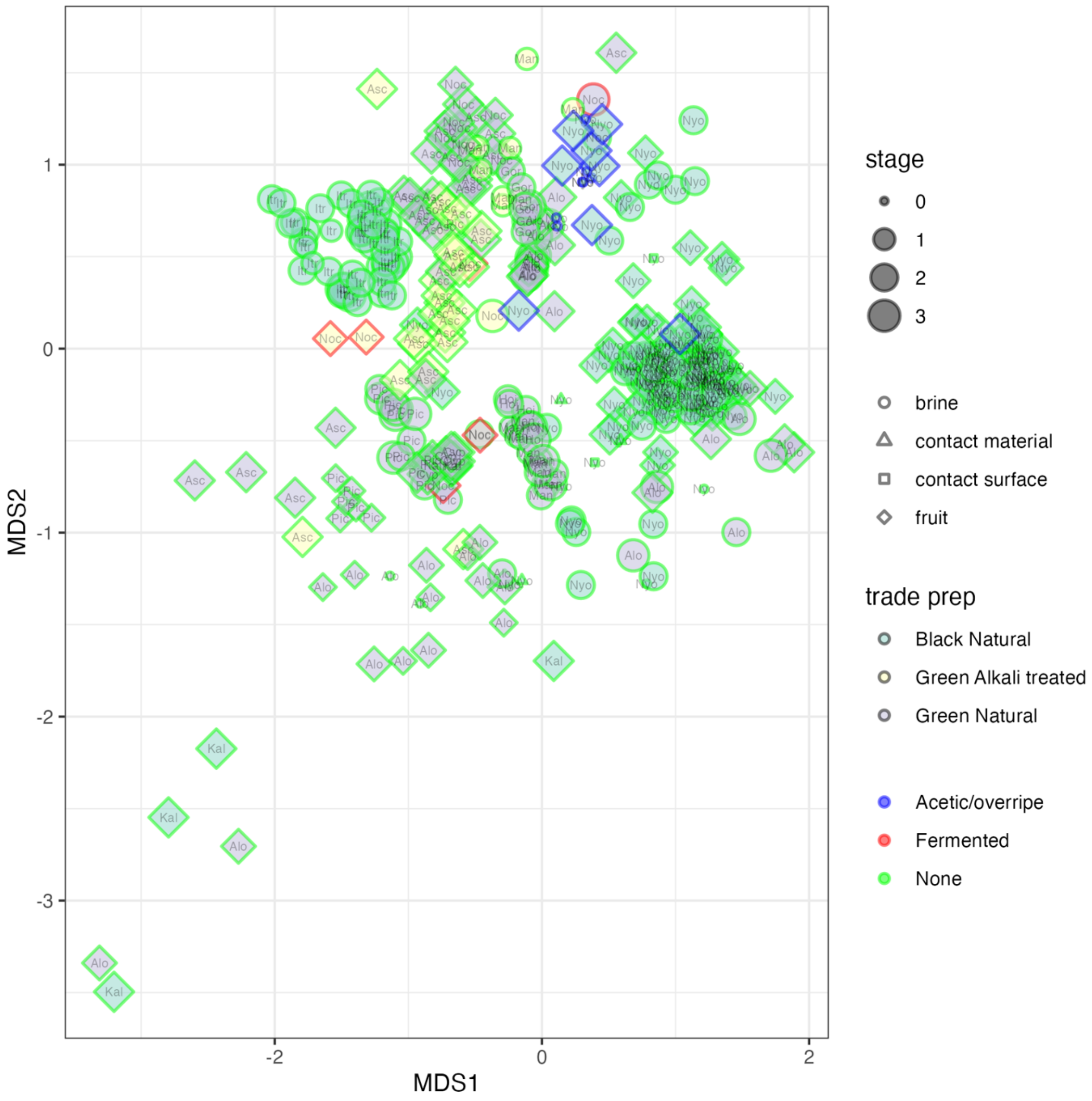


**Supplementary Figure 6**. Ordination showing the sample coordinates after non-monotonic Multidimensional Scaling of the Bray-Curtis distance matrix for the composition of bacterial microbiota of samples belonging to the METAOlive FMBN data set after prevalence and abundance filtering. Samples are divided by combination of olive ripeness and trade preparation and fermentation stage (1 beginning of fermentation, 2 samples during fermentation, 3 samples at the end of fermentation, 0 other, including environmental samples). B = brine, F = fruit. See Supplementary Table 3 for abbreviations of olive varieties. In this figure spoiled samples rather than those with starter addition are highlighted.

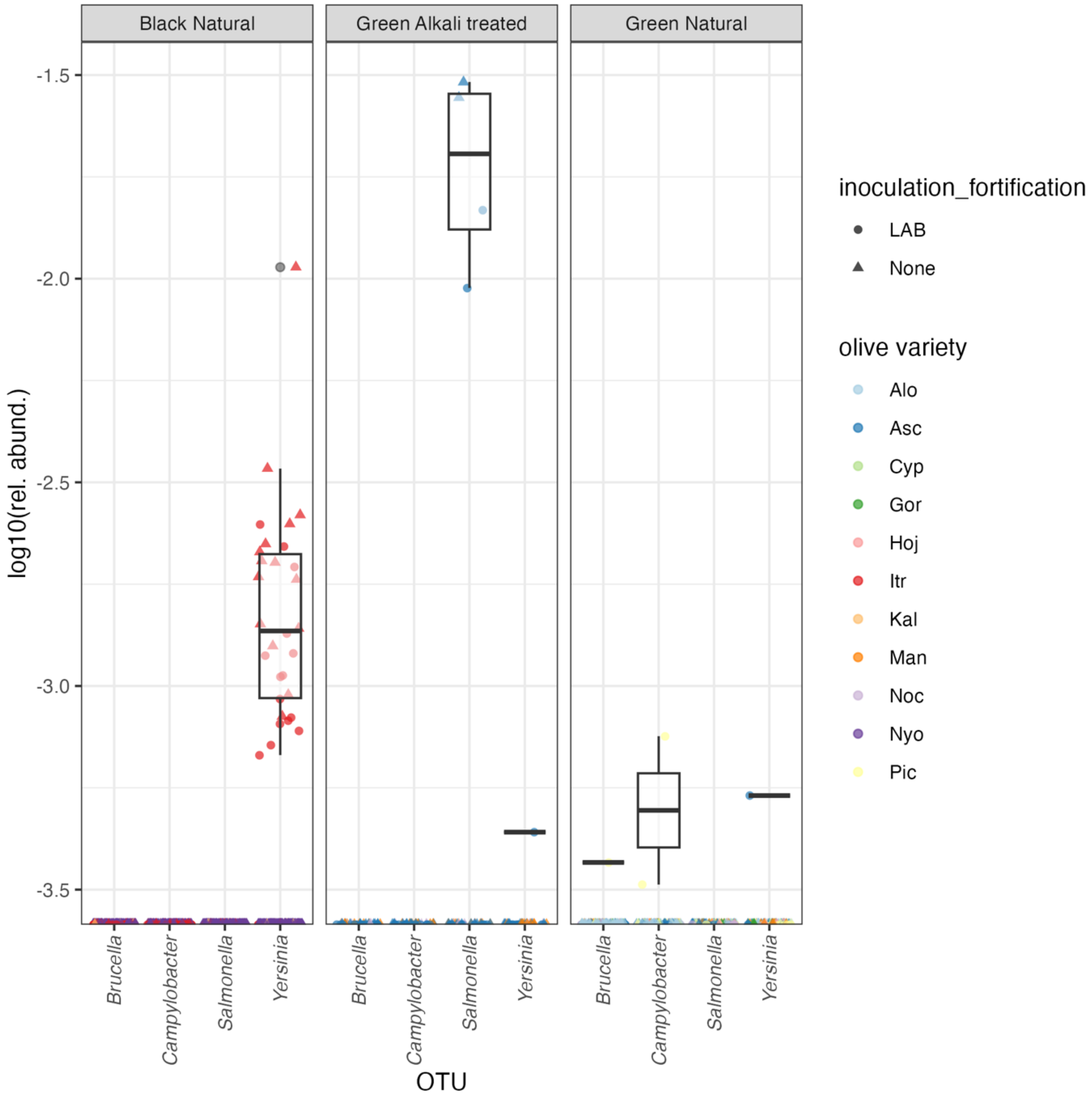


**Supplementary Figure 7**. Boxplot showing the distribution of the abundance of sequences assigned to genera including human pathogens in table olives.

**Supplementary Table 5**. Studies including data on fungi in table olives and their production environments in FoodMicrobionet 5.0 (see main text for references)

| studyId | reference | target | region | samples | year | platform |
|---|---|---|---|---|---|---|
| **ST246** | Arroyo-López et al., 2016 | ITS region | ITS1+ITS2 | 28 | 2016 | 454 GS |
| **ST239** | Penland et al., 2020 | ITS region | ITS2 | 208 | 2020 | Illumina |
| **ST182** | López-García et al., 2021 | ITS region and 16S RNA gene | ITS1 and V3-V4 | 15 | 2021 | Illumina |
| **ST247** | Penland et al., 2021 | ITS region | ITS2 | 47 | 2021 | Illumina |
| **ST248** | Benítez-Cabello et al., 2022 | ITS region | ITS1 | 69 | 2022 | Illumina |
| **ST226** | Kamilari et al., 2023 | ITS region and 16S RNA gene | ITS1 and V3-V4 | 12 | 2023 | Illumina |
| **ST236** | Ruiz-Barba et al., 2023 | ITS region and 16S RNA gene | ITS1 and V3-V4 | 17 | 2023 | Illumina |
| **ST253** | Giavcalisco et al., 2023 | ITS region and 16S RNA gene | ITS2 and V3-V4 | 49 | 2024 | Illumina |

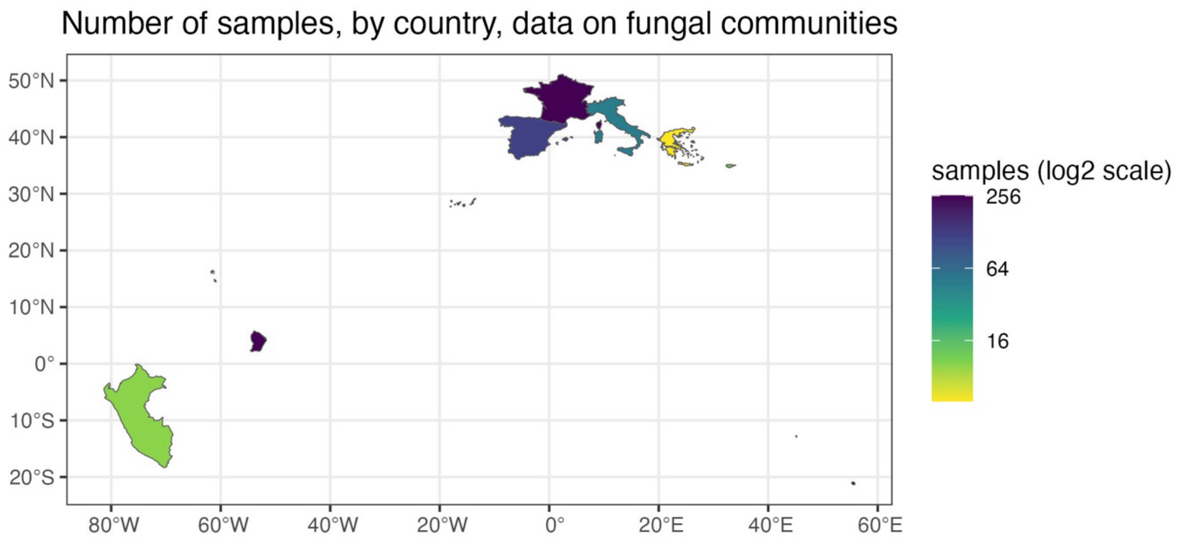


**Supplementary Figure 8**. Distribution of samples by country, data on fungal communities of table olives.

**Supplementary Table 6.** Samples with data on fungal microbiota, post sample filtering, by olive trade preparation, olive variety and sample type

| Olive trade preparation | Olive cultivar | Olive variety | Olive var. abbrev. | Olive ripeness | Sample type | n |
|---|---|---|---|---|---|---|
| **Alkali treated olives** | Conservolea | Conservolea | Con | Green olives | fruit | 3 |
| **Alkali treated olives** | Gordal | Gordal | Gor | Green olives | fruit | 2 |
| **Alkali treated olives** | Hojiblanca | Hojiblanca | Hoj | Green olives | fruit | 8 |
| **Alkali treated olives** | Lucques du Languedoc | Lucques du Languedoc | Luc | Green olives | fruit | 2 |
| **Alkali treated olives** | Manzanilla | Manzanilla | Man | Green olives | brine | 9 |
| **Alkali treated olives** | Manzanilla | Manzanilla | Man | Green olives | fruit | 12 |
| **Alkali treated olives** | Picholine | Picholine | Pic | Green olives | fruit | 2 |
| **Alkali treated olives** | Unknown | Unknown | Unk | Green olives | fruit | 2 |
| **Alkali treated olives** | Verdial | Verdial | Ver | Green olives | fruit | 2 |
| **Natural olives** | Aloreña | Aloreña | Alo | Green olives | fruit | 13 |
| **Natural olives** | Aloreña | Aloreña de Malaga | Alo | Green olives | brine | 12 |
| **Natural olives** | Aloreña | Aloreña de Malaga | Alo | Green olives | fruit | 22 |
| **Natural olives** | Arauco | Arauco | Ara | Green olives | fruit | 2 |
| **Natural olives** | Azapa | Azapa | Aza | Black olives | fruit | 2 |
| **Natural olives** | Azapa | Azapa | Aza | Green olives | fruit | 2 |
| **Natural olives** | Criolla | Criolla | Cri | Black olives | fruit | 2 |
| **Natural olives** | Criolla | Criolla | Cri | Green olives | fruit | 2 |

| | | | | | | |
|---|---|---|---|---|---|---|
| **Natural olives** | Cypriot | Cypriot | Cyp | Green olives | fruit | 6 |
| **Natural olives** | Empletre | Empletre | Emp | Green olives | fruit | 4 |
| **Natural olives** | Galega | Galega | Gal | Black olives | fruit | 2 |
| **Natural olives** | Gordal | Gordal | Gor | Green olives | brine | 6 |
| **Natural olives** | Hojiblanca | Hojiblanca | Hoj | Green olives | brine | 6 |
| **Natural olives** | Itrana | Itrana | Itr | Olives turning color | brine | 48 |
| **Natural olives** | Kalamata | Kalamata | Kal | Black olives | fruit | 5 |
| **Natural olives** | Manzanilla | Manzanilla | Man | Green olives | brine | 14 |
| **Natural olives** | Picual | Picual | Pic | Green olives | fruit | 3 |
| **Natural olives** | Tanche | Nyons | Nyo | Black olives | brine | 123 |
| **Natural olives** | Tanche | Nyons | Nyo | Black olives | contact material | 12 |
| **Natural olives** | Tanche | Nyons | Nyo | Black olives | contact surface | 7 |
| **Natural olives** | Tanche | Nyons | Nyo | Black olives | fruit | 113 |
| **Natural olives** | Unknown | Unknown | Unk | Black olives | fruit | 2 |
| **Natural olives** | Unknown | Unknown | Unk | Green olives | fruit | 2 |
| **Unknown** | Manzanilla | Manzanilla | Man | Green olives | fruit | 1 |

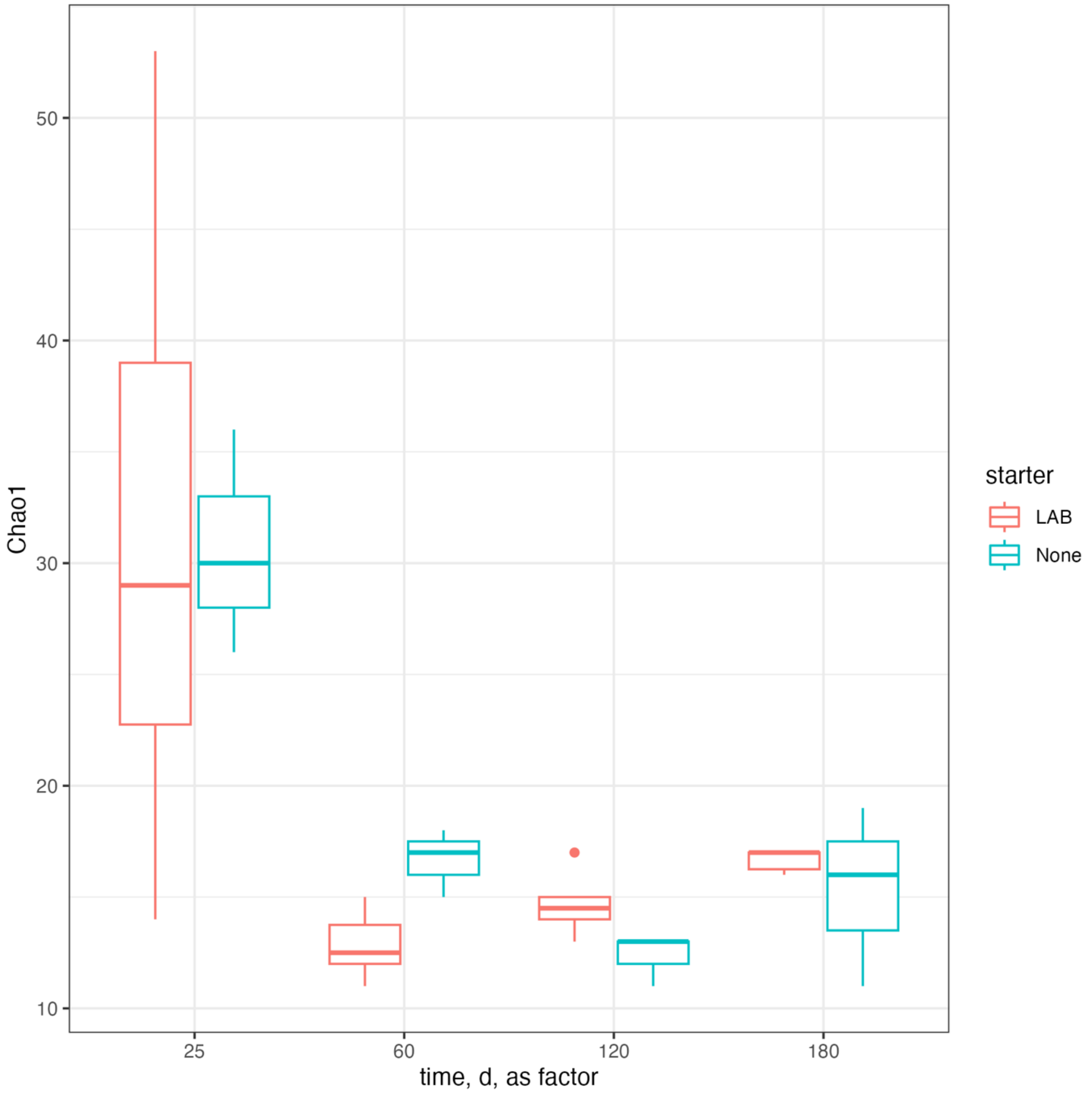


**Supplementary Figure 9.** Time course of diversity of fungal communities in brines of started (*Lactipl. pentosus*) and non-started Itrana olives.

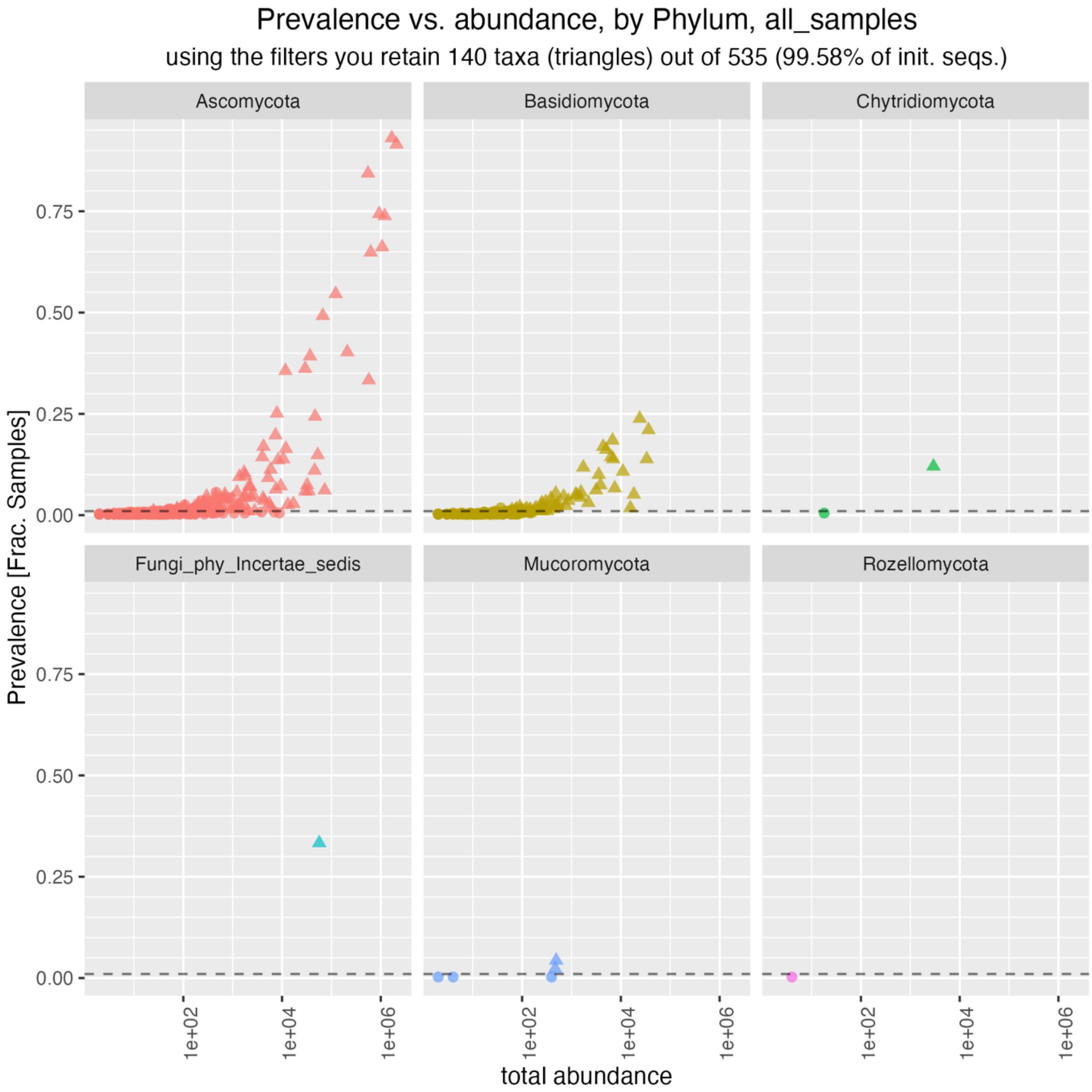


**Supplementary Figure 10.** A prevalence and abundance plot for fungal genera in table olives, METAOlive dataset.

**Supplementary Table 7**. Prevalence and abundance data for the core fungal genera in table olives.

| Genus | Order | Prevalence | Total Ab. | Min. rel. ab. | Max. rel. ab. | Mean. rel. ab. | Rel. prev. |
|---|---|---|---|---|---|---|---|
| ***Candida*** | Saccharomycetales | 357 | 2084122 | 0 | 1.00 | 2.09E-01 | 0.92 |
| ***Pichia*** | Saccharomycetales | 363 | 1668725 | 0 | 1.00 | 1.67E-01 | 0.93 |
| ***Saccharomyces*** | Saccharomycetales | 288 | 1199799 | 0 | 0.95 | 1.20E-01 | 0.74 |
| ***Citeromyces*** | Saccharomycetales | 258 | 1065449 | 0 | 0.81 | 1.07E-01 | 0.66 |
| ***Zygotorulaspora*** | Saccharomycetales | 290 | 927747 | 0 | 0.89 | 9.31E-02 | 0.74 |
| ***Wickerhamomyces*** | Saccharomycetales | 329 | 551146 | 0 | 0.77 | 5.53E-02 | 0.84 |
| ***Aureobasidium*** | Dothideales | 253 | 624762 | 0 | 0.99 | 6.27E-02 | 0.65 |
| ***Nakazawaea*** | Saccharomycetales | 130 | 572201 | 0 | 0.66 | 5.74E-02 | 0.33 |
| ***Cladosporium*** | Cladosporiales | 213 | 121763 | 0 | 1.00 | 1.22E-02 | 0.55 |
| ***Alternaria*** | Pleosporales | 157 | 209870 | 0 | 0.97 | 2.11E-02 | 0.40 |
| ***Penicillium*** | Eurotiales | 192 | 66812 | 0 | 0.66 | 6.70E-03 | 0.49 |
| ***Phaeococcomyces*** | Lichenostigmatales | 153 | 36859 | 0 | 0.18 | 3.70E-03 | 0.39 |
| ***Zygoascus*** | Saccharomycetales | 141 | 29404 | 0 | 0.04 | 2.95E-03 | 0.36 |
| ***Fungi gen Incertae sedis*** | Fungi gen Incertae sedis | 130 | 56657 | 0 | 0.93 | 5.68E-03 | 0.33 |
| ***Priceomyces*** | Saccharomycetales | 139 | 11702 | 0 | 0.03 | 1.17E-03 | 0.36 |
| ***Wickerhamiella*** | Saccharomycetales | 95 | 46604 | 0 | 0.81 | 4.68E-03 | 0.24 |
| ***Schwanniomyces*** | Saccharomycetales | 98 | 7846 | 0 | 0.06 | 7.87E-04 | 0.25 |
| ***Filobasidium*** | Filobasidiales | 93 | 24069 | 0 | 0.39 | 2.41E-03 | 0.24 |
| ***Vishniacozyma*** | Tremellales | 82 | 36266 | 0 | 0.56 | 3.64E-03 | 0.21 |
| ***Coniozyma*** | Dothideales | 77 | 7388 | 0 | 0.06 | 7.41E-04 | 0.20 |
| ***Symmetrospora*** | Cystobasidiomycetes ord Incertae sedis | 72 | 6781 | 0 | 0.04 | 6.80E-04 | 0.18 |
| ***Starmerella*** | Saccharomycetales | 58 | 52665 | 0 | 0.94 | 5.28E-03 | 0.15 |
| ***Geotrichum*** | Saccharomycetales | 64 | 11917 | 0 | 0.72 | 1.20E-03 | 0.16 |
| ***Sporobolomyces*** | Sporidiobolales | 66 | 4378 | 0 | 0.06 | 4.39E-04 | 0.17 |
| ***Neoascochyta*** | Pleosporales | 66 | 4209 | 0 | 0.20 | 4.22E-04 | 0.17 |
| ***Malassezia*** | Malasseziales | 63 | 5044 | 0 | 0.16 | 5.06E-04 | 0.16 |
| ***Kwoniella*** | Tremellales | 54 | 33331 | 0 | 0.11 | 3.34E-03 | 0.14 |
| ***Rhodotorula*** | Sporidiobolales | 56 | 6334 | 0 | 0.20 | 6.36E-04 | 0.14 |
| ***Saccharomycopsis*** | Saccharomycetales | 56 | 3988 | 0 | 0.04 | 4.00E-04 | 0.14 |
| ***Debaryomyces*** | Saccharomycetales | 54 | 10595 | 0 | 0.10 | 1.06E-03 | 0.14 |
| ***Cystofilobasidium*** | Cystofilobasidiales | 54 | 7000 | 0 | 0.09 | 7.02E-04 | 0.14 |
| ***Aspergillus*** | Eurotiales | 43 | 45703 | 0 | 0.89 | 4.59E-03 | 0.11 |
| ***Yamadazyma*** | Saccharomycetales | 53 | 8286 | 0 | 0.21 | 8.31E-04 | 0.14 |
| ***Chytridiomycota gen Incertae sedis*** | Chytridiomycota gen Incertae sedis | 47 | 2945 | 0 | 0.54 | 2.95E-04 | 0.12 |
| ***Dioszegia*** | Tremellales | 46 | 1733 | 0 | 0.07 | 1.74E-04 | 0.12 |
| ***Myriangiales gen Incertae sedis*** | Myriangiales | 44 | 5873 | 0 | 0.10 | 5.89E-04 | 0.11 |
| ***Naganishia*** | Filobasidiales | 42 | 11009 | 0 | 0.14 | 1.10E-03 | 0.11 |

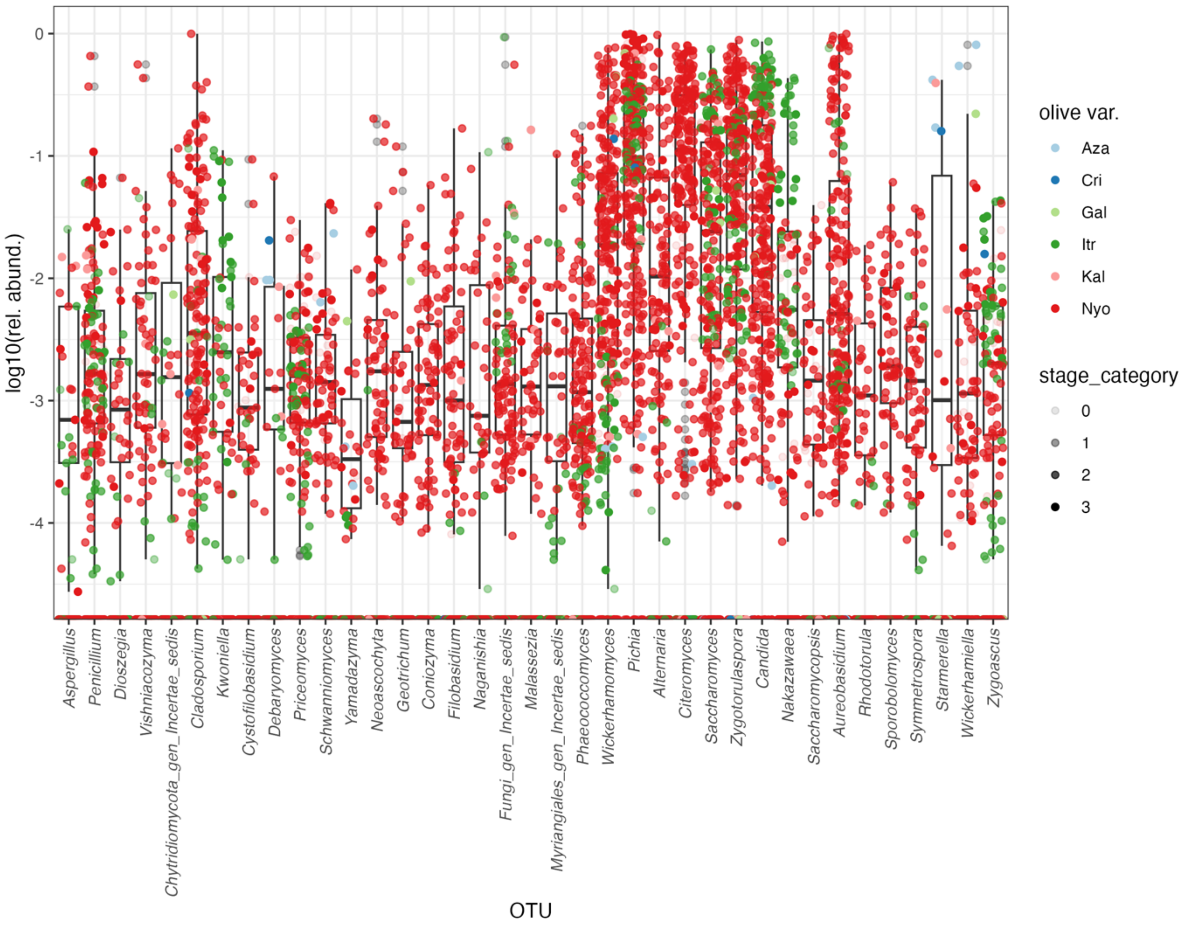


**Supplementary Figure 11**. Distribution of the dominating fungal genera in black natural olives. Refer to Supplementary table 3 for olive variety abbreviations.

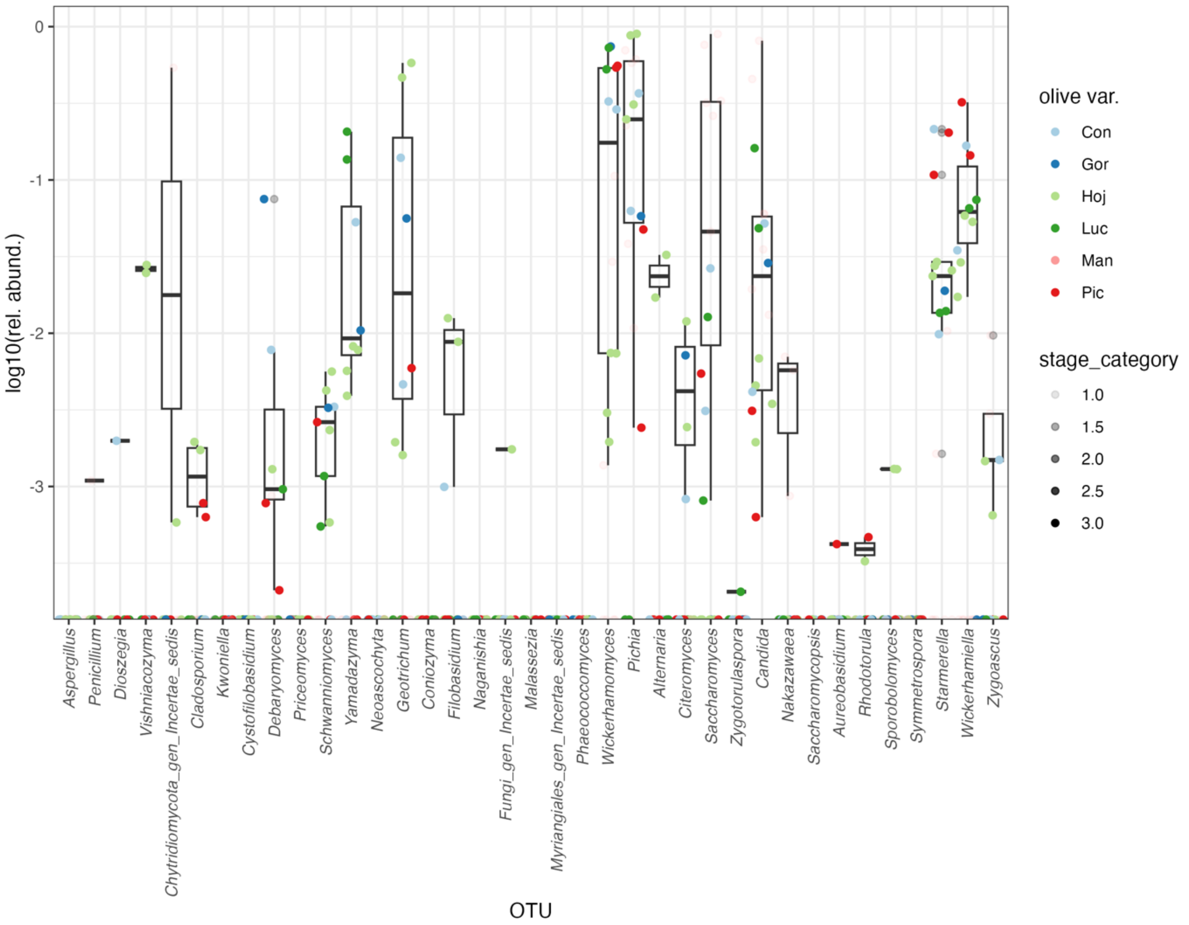


**Supplementary Figure 12**. Distribution of the dominating fungal genera in green alkali treated olives. Refer to Supplementary table 3 for olive variety abbreviations.

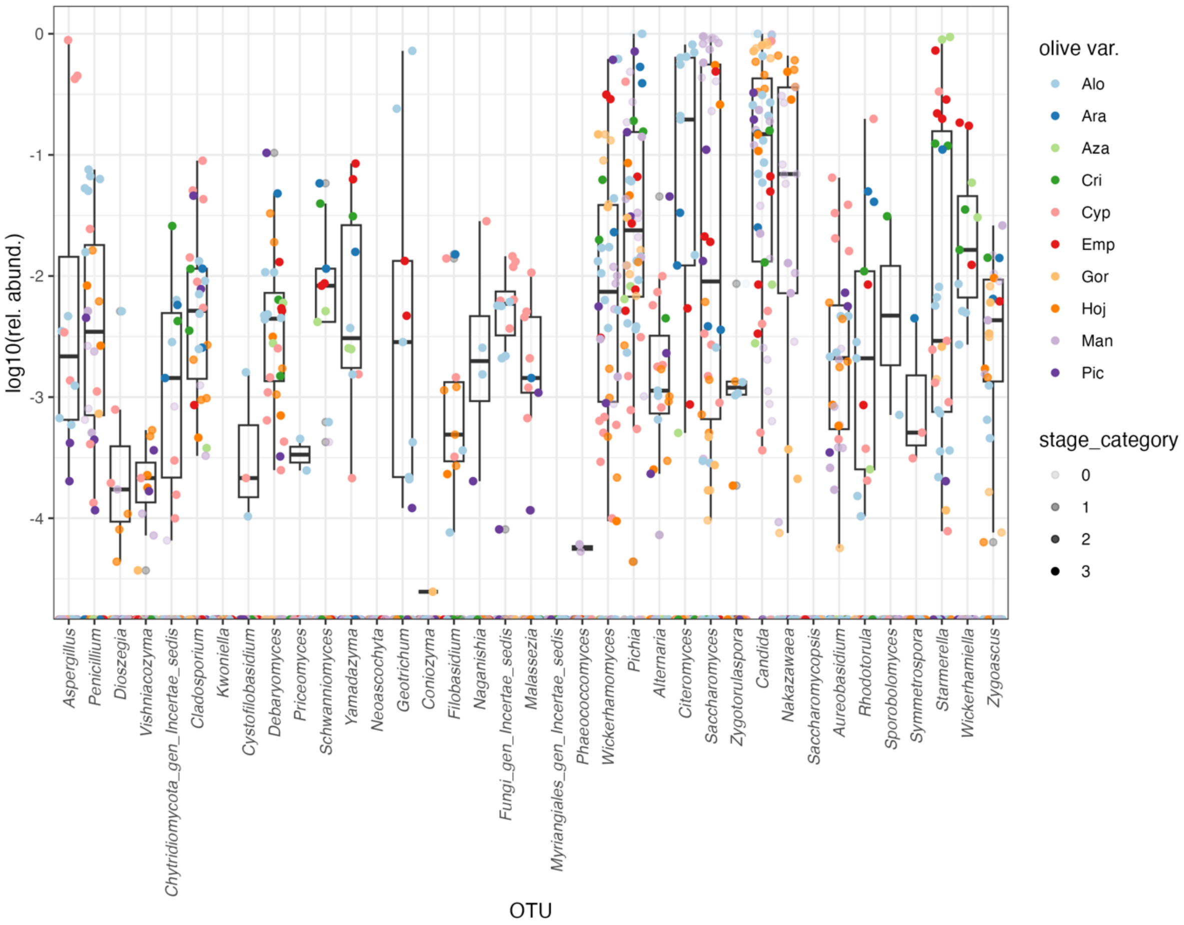


**Supplementary Figure 13**. Distribution of the dominating fungal genera in Green natural olives. Refer to Supplementary Table 3 for olive variety abbreviations.

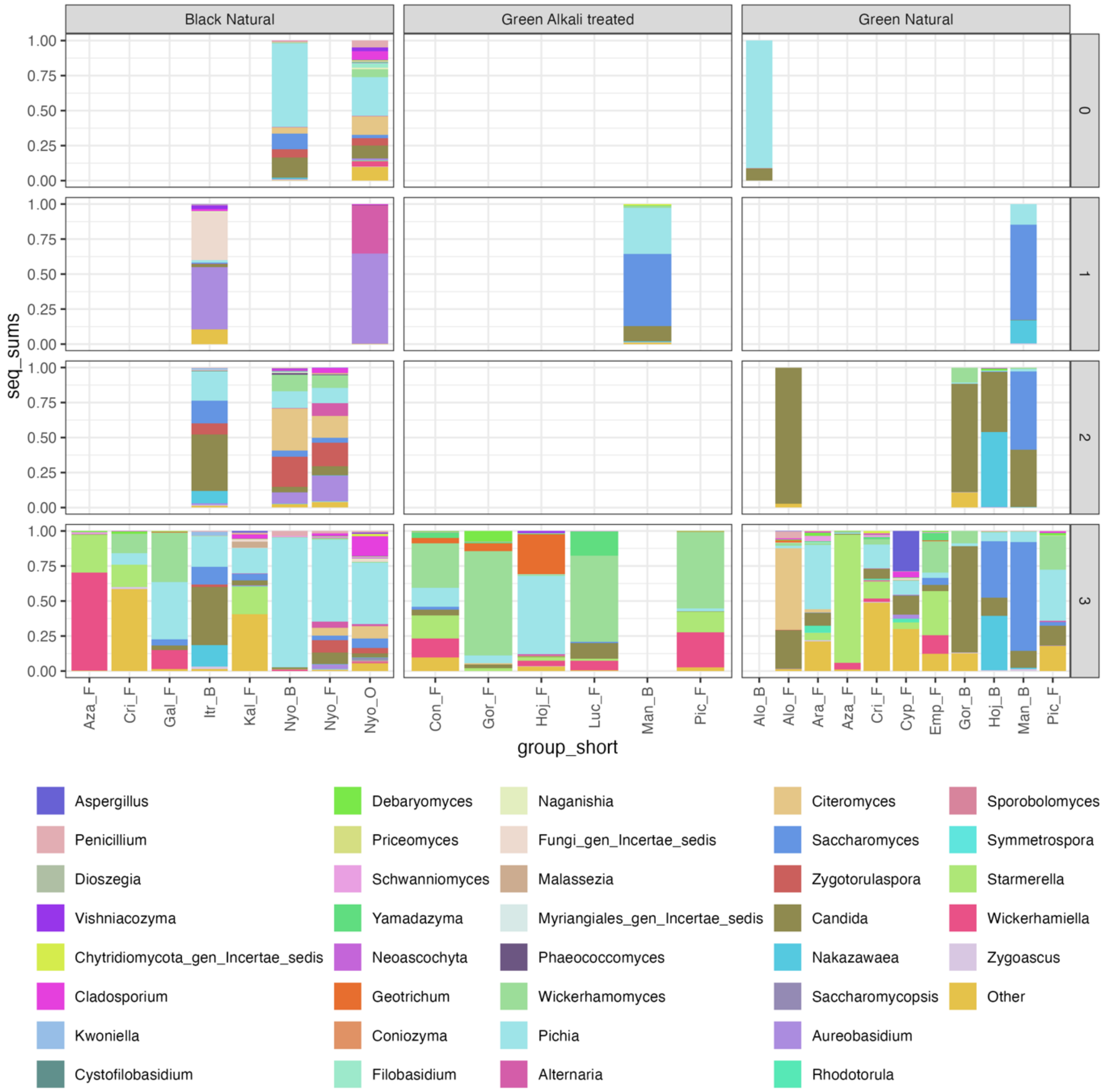


**Supplementary Figure 14**. Composition of fungal communities of different varieties of table olives and fermentation brine in the METAOlive FMBN data set. The most prevalent and abundant genera are shown; the others are pooled under the group “Other”. Samples are divided by combination of olive ripeness and trade preparation and fermentation stage (1 beginning of fermentation, 2 samples during fermentation, 3 samples at the end of fermentation, 0 other). B = brine, F = fruit. See Supplementary Table 3 for abbreviations of olive varieties.

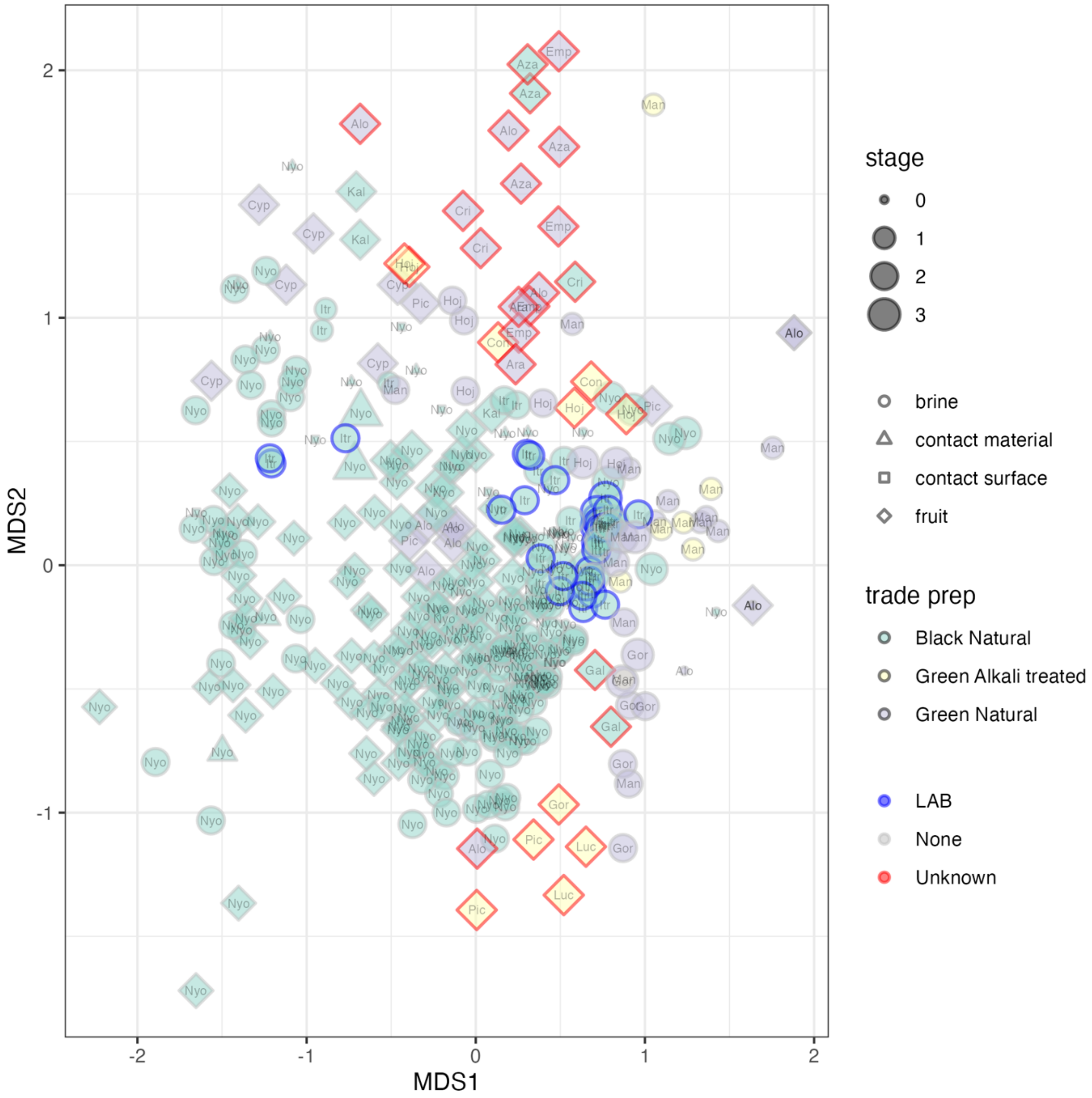


**Supplementary Figure 15**. Ordination showing the sample coordinates after non-monotonic Multidimensional Scaling of the Bray-Curtis distance matrix for the composition of fungal microbiota of samples belonging to the METAOlive FMBN data set after prevalence and abundance filtering. Samples are divided by combination of olive ripeness and trade preparation and fermentation stage (1 beginning of fermentation, 2 samples during fermentation, 3 samples at the end of fermentation, 0 other, including environmental samples). B = brine, F = fruit. See Supplementary Table 3 for abbreviations of olive varieties.

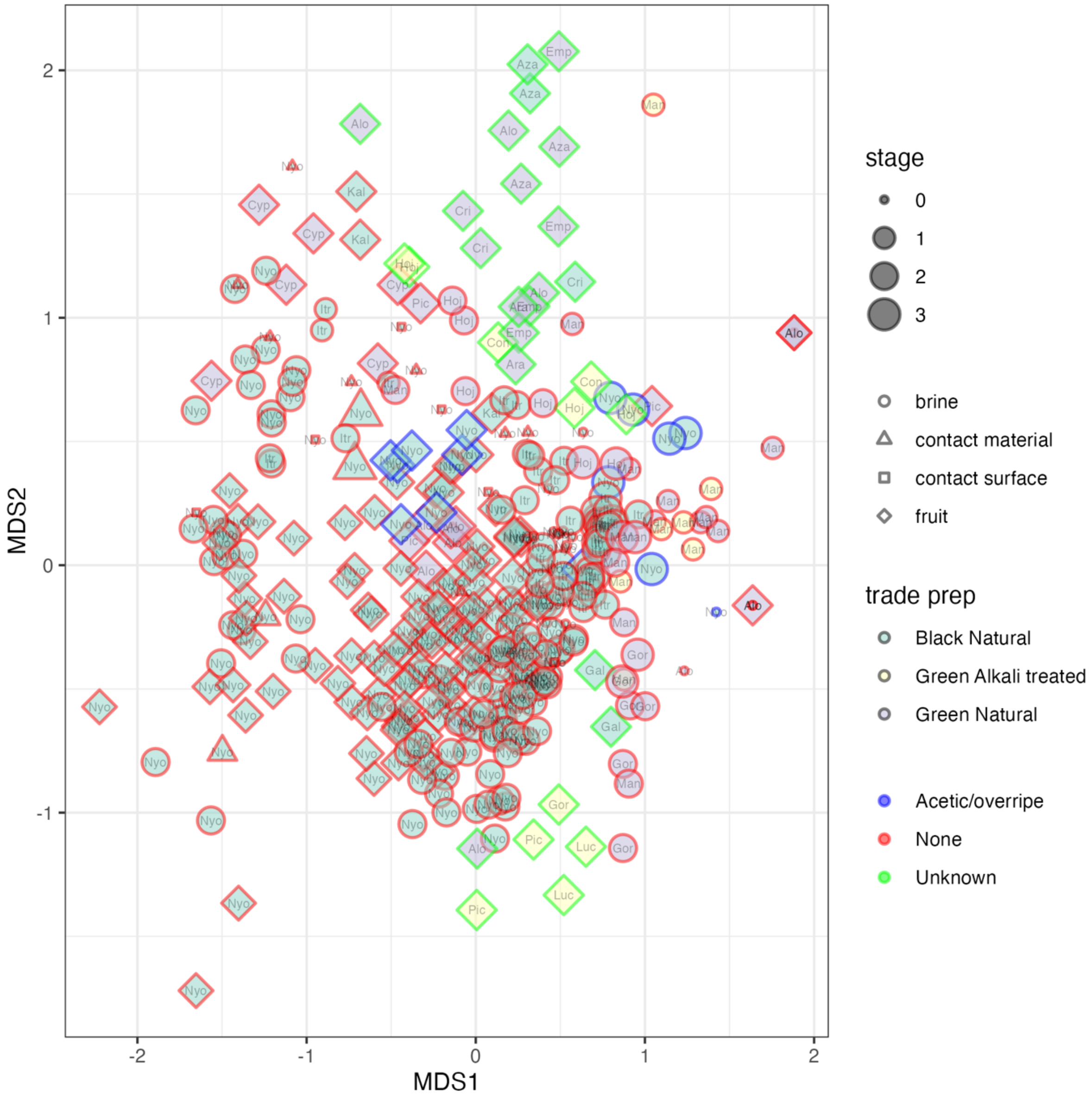


**Supplementary Figure 16**. Ordination showing the sample coordinates after non-monotonic Multidimensional Scaling of the Bray-Curtis distance matrix for the composition of fungal microbiota of samples belonging to the METAOlive FMBN data set after prevalence and abundance filtering. Samples are divided by combination of olive ripeness and trade preparation and fermentation stage (1 beginning of fermentation, 2 samples during fermentation, 3 samples at the end of fermentation, 0 other, including environmental samples). B = brine, F = fruit. Information on spoilage is included. See Supplementary Table 3 for abbreviations of olive varieties.
